# CXCR3-expressing myeloid cells recruited to the hypothalamus protect against diet-induced body mass gain and metabolic dysfunction

**DOI:** 10.1101/2024.01.16.575893

**Authors:** Natalia F. Mendes, Ariane M. Zanesco, Cristhiane F. Aguiar, Gabriela F. Rodrigues-Luiz, Dayana C. da Silva, Jonathan F. Campos, Niels O. S. Câmara, Pedro M. M. de Moraes-Vieira, Eliana P. de Araújo, Licio A. Velloso

**Affiliations:** School of Medical Sciences, Department of Translational Medicine (Section of Pharmacology), University of Campinas, Brazil; Laboratory of Cell Signaling, Obesity and Comorbidities Research Center, University of Campinas, Brazil; Laboratory of Immunometabolism, Institute of Biology - University of Campinas, Brazil; Department of Microbiology, Immunology and Parasitology, Federal University of Santa Catarina, Brazil; Laboratory for Transplantation Immunobiology, Institute of Biomedical Sciences, University of Sao Paulo, Brazil; Faculty of Nursing, University of Campinas, Brazil; National Institute of Science and Technology on Neuroimmunomodulation, Rio de Janeiro, Brazil

**Keywords:** Hypothalamus, obesity, monocytes, microglia, chemokines

## Abstract

Microgliosis is an important component of diet-induced hypothalamic inflammation in obesity. A few hours after the introduction of a high-fat diet, the mediobasal hypothalamus resident microglia undergo morphological and functional changes toward an inflammatory phenotype. If the consumption of large amounts of dietary fats persists for long periods, bone marrow-derived myeloid cells are recruited and integrated into a new landscape of hypothalamic microglia. However, it is currently unknown what are the transcriptional signatures and specific functions exerted by either resident or recruited subsets of hypothalamic microglia. Here, the elucidation of the transcriptional signatures revealed that resident microglia undergo only minor changes in response to dietary fats; however, under the consumption of a high-fat diet, there are major transcriptional differences between resident and recruited immune cells with major impact on chemotaxis. In addition, in CCR2+ recruited peripheral immune cells, there are major transcriptional differences between females and males with important impact on transcripts involved in neurodegeneration and thermogenesis. The chemokine receptor CXCR3 emerged as one of the components of chemotaxis with the greatest difference between recruited and resident microglia, and thus, was elected for further intervention. The hypothalamic immunoneutralization of CXCL10, one of the ligands for CXCR3, resulted in increased body mass gain and reduced energy expenditure, particularly in females. Furthermore, the chemical inhibition of CXCR3 resulted in a much greater change in phenotype with increased body mass gain, reduced energy expenditure, increased blood leptin, glucose intolerance, and reduced insulin. Thus, this study has elucidated the transcriptional differences between resident microglia and recruited immune cells in diet-induced obesity, identifying chemokines as a relevant subset of genes undergoing regulation. In addition, we showed that a subset of recruited immune cells expressing CXCR3 has a protective, rather than a detrimental role in the metabolic outcomes promoted by the consumption of a high-fat diet, thus, establishing a new concept in obesity-associated hypothalamic inflammation.

## Introduction

Obesity affects over 600 million people worldwide and projections are pessimistic indicating that over a billion people will be diagnosed with obesity by the year 2030 (worldobesity.org). Obesity develops as a consequence of a chronic state of anabolism in which caloric intake overcomes energy expenditure (Theilade et al., 2021). Both, experimental and human studies have indicated that, at least in part, the chronic anabolic state leading to obesity develops as a consequence of a defective hypothalamic regulation of whole-body energy balance (Cavadas et al., 2016) (Sonnefeld et al., 2023) (van de Sande-Lee et al., 2020).

Experimental studies have shown that the consumption of large amounts of saturated fats triggers an inflammatory response in the hypothalamus affecting the function of key neurons involved in the regulation of food intake, energy expenditure, and systemic metabolism (De Souza et al., 2005) (Milanski et al., 2009) (Zhang et al., 2008) (Milanski et al., 2012). In addition, human studies using magnetic resonance imaging have identified hypothalamic gliosis in adults and children with obesity, providing clinical evidence for the existence of an obesity-associated hypothalamic inflammation (van de Sande-Lee et al., 2020) (Thaler et al., 2012) (Sewaybricker et al., 2019). Microglia are key cellular components of this inflammatory response undergoing structural and functional changes that develop early after the introduction of a high-fat diet (HFD) (Tapia-González et al., 2011) (Valdearcos et al., 2014) (Fernández-Arjona et al., 2022) (Valdearcos et al., 2017). Distinct strategies used to inhibit hypothalamic microglia resulted in impaired diet-induced hypothalamic inflammation, increased leptin sensitivity, reduced spontaneous caloric intake, and improved systemic glucose tolerance (Valdearcos et al., 2014) (Morari et al., 2014). Thus, elucidating the mechanisms involved in the regulation of microglial response to dietary factors may provide advance in the definition of the pathophysiology of obesity, and potentially identify new targets for interventions aimed at treating obesity and its metabolic comorbidities (Mendes et al., 2018) (Mendes and Velloso, 2022) (Valdearcos et al., 2019).

Resident microglia are derived from the yolk sac primitive hematopoietic cells and populate the neuroepithelium at embryonic day 9.5 (Ginhoux et al., 2010). Under baseline conditions, in the absence of infections, trauma or other types of potentially harmful stimuli, resident microglia remain rather quiescent; however, under stimulus, they undergo rapid morphological and functional changes aimed at confronting the threat (Saijo and Glass, 2011). In the hypothalamus, differently from most parts of the brain, resident microglia are responsive to fluctuations in the blood levels of nutrients and hormones involved in metabolic control, such as leptin (Valdearcos et al., 2014). Thus, a simple meal can promote considerable changes in hypothalamic resident microglia indicating the involvement of these cells in the complex network of cells that regulate whole-body metabolism (Gao et al., 2014). Under the consumption of a nutritionally balanced diet, meal-induced changes in hypothalamic resident microglia are cyclic and completely reversible (Gao et al., 2014). However, under consumption of a HFD, microglia present profound changes in morphology and function; moreover, upon prolonged consumption of this type of diet, there is recruitment of bone marrow-derived myeloid cells that will compose a new landscape of hypothalamic microglia (Valdearcos et al., 2014) (Morari et al., 2014) (Valdearcos et al., 2019). Despite the considerable advance in the understanding of how hypothalamic microglia are involved in diet-induced obesity (DIO), it is currently unknown what are the transcriptional landscapes of resident microglia and recruited immune cells and what specific functions either subset of microglia exerts in the context of obesity and metabolic control.

In this study, we first elucidated the transcriptional differences between resident CX3CR1+ microglial cells and CCR2+ recruited peripheral immune cells in DIO, identifying chemokines as a relevant subset of genes undergoing regulation. Next, we identified a subset of recruited microglia expressing CXCR3, which exerts a protective role against DIO.

## Results

### Elucidating the transcriptional signatures of hypothalamic resident microglia and recruited immune cells in diet-induced obesity

Double reporter mice were obtained by the crossing of CX3CR1^GFP^ and CCR2^RFP^ (Fig. 1a). At the age of 8 weeks, female and male mice were randomly selected for either chow or HFD feeding for 28 days, and then specimens were harvested for analysis (Fig. 1b). In flow cytometry, cells expressing CCR2 could not be detected in the hypothalamus of mice fed on chow (Fig. 1c), whereas in the white adipose tissue, virtually all cells expressing CX3CR1 were also expressing CCR2 (Fig. 1c). Conversely, in both female and male mice fed on HFD, CCR2 cells accounted for approximately 10% of hypothalamic microglia (Fig. 1d). Histological analysis confirmed the results obtained by flow cytometry (Fig. 1e); furthermore, it was shown that CD169, which is classically regarded as a marker of bone marrow-derived cells (Chávez-Galán et al., 2015), is in fact expressed both in resident microglia as well as recruited immune cells (Fig. 1f), confirming data published elsewhere (Valdearcos et al., 2019). Next, to prepare the samples for RNA sequencing, we sorted hypothalamic microglia expressing either CX3CR1 and recruited cells expressing CCR2 (Fig. 2a). The quality of sorting was confirmed by determining the positivity for CX3CR1 (Fig. 2b) and CCR2 (Fig. 2c), and by determining the positivity for several markers of resident microglia (Fig. 2d-2m) and several markers of bone marrow-derived cells (Fig. 2n-2x). The elucidation of the transcriptional landscapes of CX3CR1 and CCR2 cells was performed using bioinformatic tools to compare transcript expression levels in distinct cell types and conditions. The results (Table 1) revealed that either diet or sex exerted only minor differences in the expression of transcripts in CX3CR1 cells (Fig. 3a-3b); nevertheless, sex exerted major differences in CCR2 cells (Fig. 3a-3b). The direct comparisons between CX3CR1 and CCR2 obtained from mice fed on the HFD, revealed the vast differences in the transcriptional landscapes of either female (Fig. 3c) or male (Fig. 3d) mice. Furthermore, the direct comparisons between female and male mice revealed a considerable degree of sexual dimorphism in the transcriptional landscapes of recruited CCR2 cells (Fig. 3e). In CX3CR1 cells, the consumption of the HFD impacted on IL17, lipids, toll-like receptor signaling, tumor necrosis factor signaling and chemokines (Fig. 4a); whereas in CCR2 cells, the consumption of the HFD impacted on lipids, toll-like receptor signaling, tumor necrosis factor signaling, chemokines, neurotrophins signaling, reactive oxygen species, thermogenesis, and pathways related to neurodegeneration (Fig. 4a). Next, we asked what functions related to chemotaxis were predominantly regulated in CCR2 cells of mice fed on the HFD (Fig. 4b-4c). As cell chemotaxis emerged as an important function in both females (Fig. 4b) and males (Fig. 4c), we looked with greater detail into the expression of chemokines and chemokine receptors (Fig. 4d). Virtually all the transcripts evaluated showed diametrically opposite expression between CX3CR1 and CCR2 (Fig. 4d).

**Figure 1.**
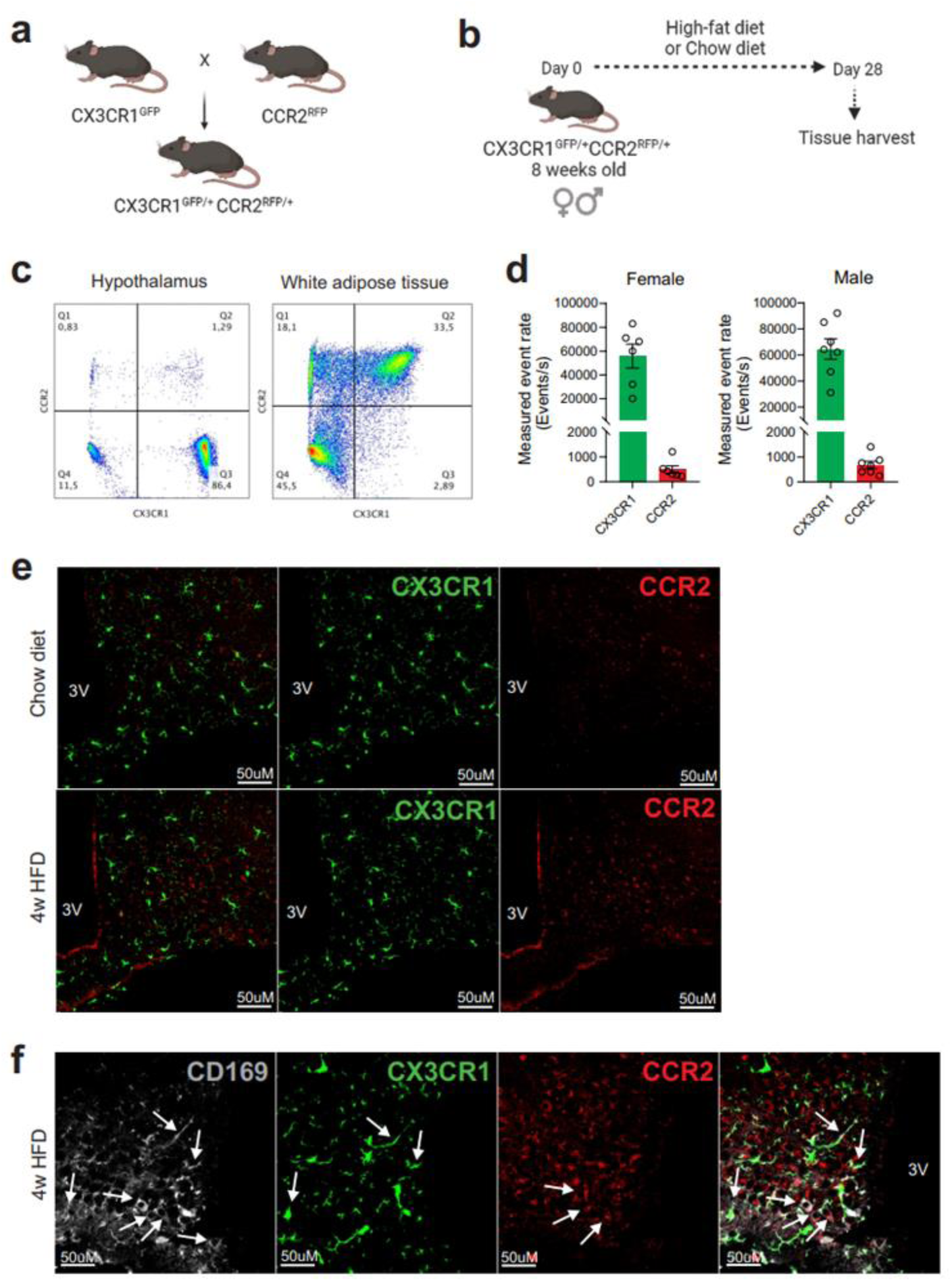
CCR2+ cells infiltrate the hypothalamus of mice fed a high-fat diet. a) CX3CR1^GFP/+^CCR2^RFP/+^ dual-reporter mutant mice generation. b) Schematic representation of the experimental protocol for analysis of HFD-induced CCR2+ peripheral-cell chemotaxis towards the hypothalamus. c) Flow cytometry analysis of CX3CR1^GFP+^ and CCR2^RFP+^ cells in the white adipose tissue and in the hypothalamus of HFD-fed mice. d) Measured event rate detected by flow cytometer of CX3CR1^GFP+^ and CCR2^RFP+^ cells isolated from the hypothalamus of HFD-fed male and female mice. e) Coronal brain sections of mediobasal hypothalamus (MBH) from chow- and 4-week HFD-fed mice CX3CR1^GFP/+^CCR2^RFP/+^. f) Coronal brain sections of MBH from 4 weeks HFD-fed mice CX3CR1^GFP/+^CCR2^RFP/+^ immunostained for CD169 (Sialoadhesin). White arrows indicate overlap between CD169+ cells with CX3CR1+ cells or with CCR2+ cells. 3V = Third ventricle, Scale Bar = 50 μm.

**Figure 2.**
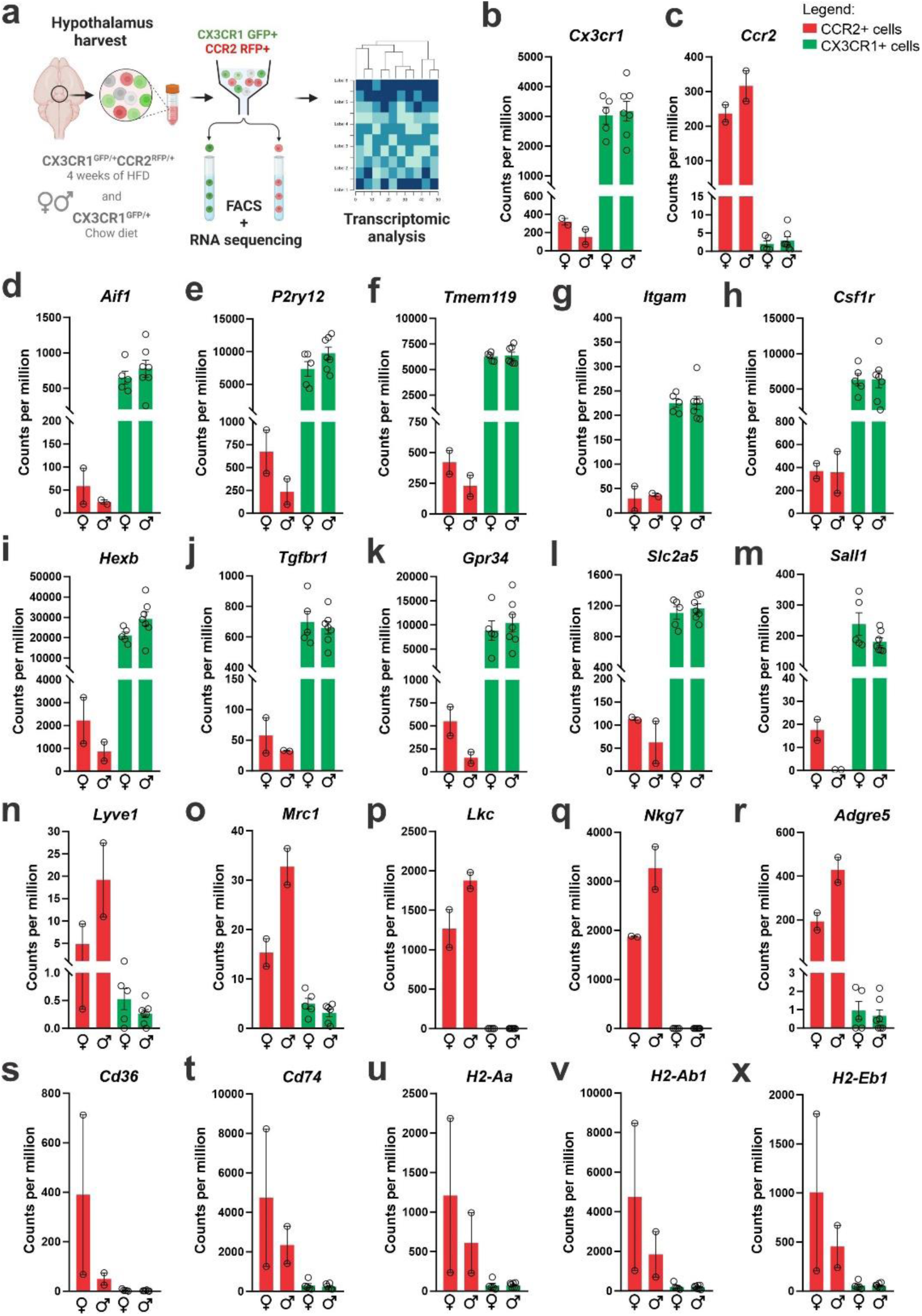
CX3CR1+ resident and CCR2+ recruited cells sorted from the hypothalamus of HFD-fed mice express classical markers of microglia and other immune cells. a) Schematic representation of the experimental protocol for sorting and sequencing CX3CR1+ and CCR2+ cells from the hypothalamus of chow- and HFD-fed mice. b) Cx3cr1 gene expression and c) Ccr2 gene expression of CX3CR1+ and CCR2+ cells sorted from the hypothalamus of HFD-fed mice. Analysis of d-m) classical microglial markers, and n-x) bone marrow-derived immune cell markers in the transcriptome of CX3CR1+ and CCR2+ cells sorted from the hypothalamus of HFD-fed mice. To perform RNA-sequencing we have employed a total of 200 mice of each sex fed on HFD and 100 mice of each sex fed on chow diet. They were divided into 5 independent experiments. To get the total RNA amount from CCR2+ cells in the hypothalamus of HFD- fed mice that was enough for library construction and RNA-seq, CCR2+ samples were pooled together in 3 samples, but only 2 samples could be sequenced due to the final RNA integrity and amount.

**Figure 3.**
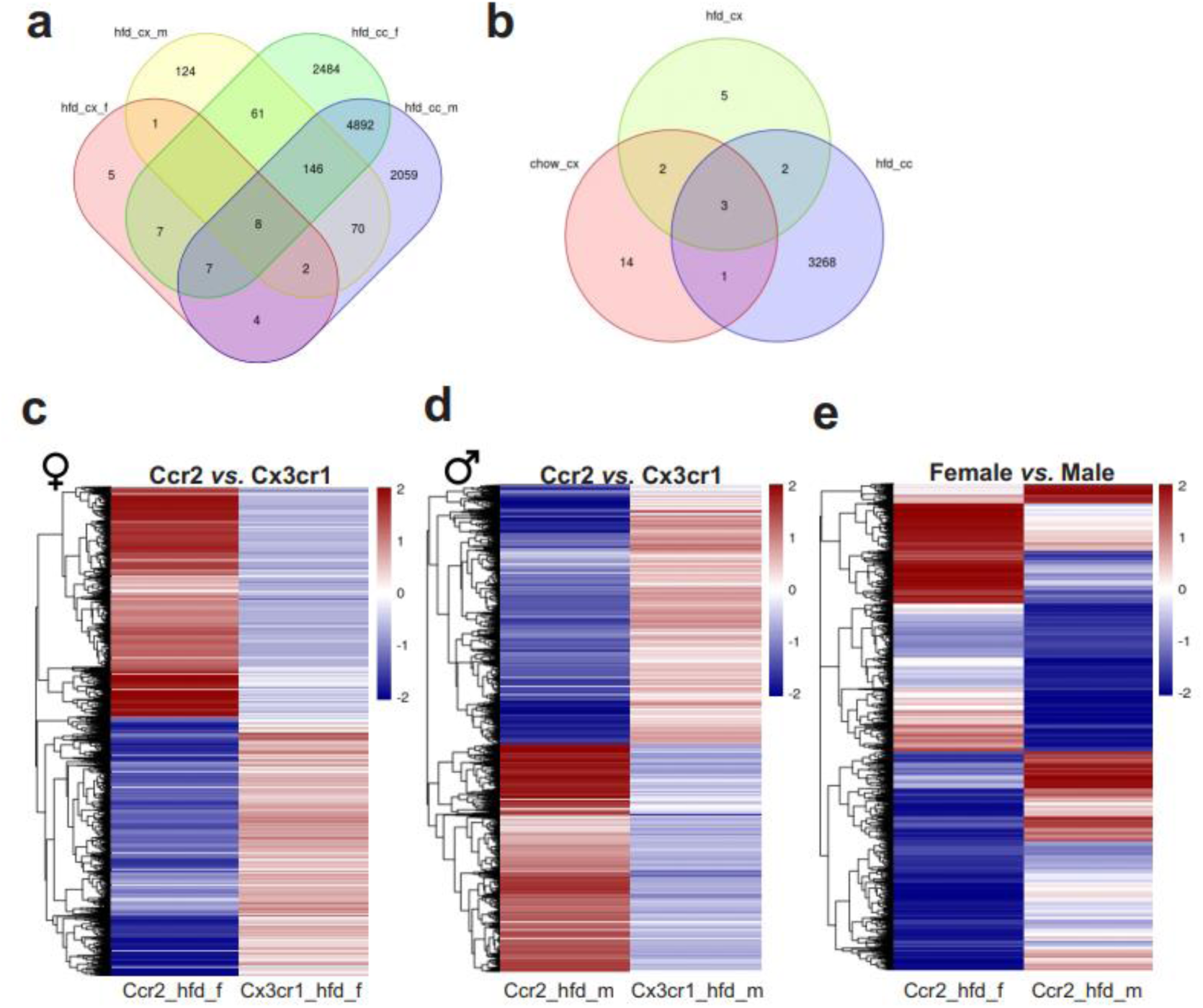
Differential gene expression (DGE) analysis of CX3CR1+ resident microglia and CCR2+ infiltrating cells sorted from the hypothalamus of HFD-fed mice show an enormous difference in their transcriptomic signature. A-b) Venn diagram showing the number of DGEs for chow and HFD diet comparison and sex comparison, respectively. c) Heatmap of up and downregulated DEGs when comparing CX3CR1+ resident and CCR2+ infiltrating cells from HFD-fed female mice. d) Heatmap of up and downregulated DEGs when comparing CX3CR1+ resident and CCR2+ infiltrating cells from HFD-fed male mice. e) Heatmap of up and downregulated DEGs when comparing CCR2+ infiltrating cells from HFD-fed male and female mice. hfd_cx_f = Cx3cr1_hfd_female vs. Cx3cr1_chow_female; hfd_cx_m = Cx3cr1_hfd_male vs. Cx3cr1_chow_male; chow_cx = Cx3cr1_chow_male vs. Cx3cr1_chow_female; hfd_cx = Cx3cr1_hfd_male vs. Cx3cr1_hfd_female; hfd_cc_f = Cx3cr1_hfd_female vs. Ccr2_hfd_female; hfd_cc_m = Cx3cr1_hfd_male vs. Ccr2_hfd_male; hfd_cc = Ccr2_hfd_male vs. Ccr2_hfd_female.

**Figure 4.**
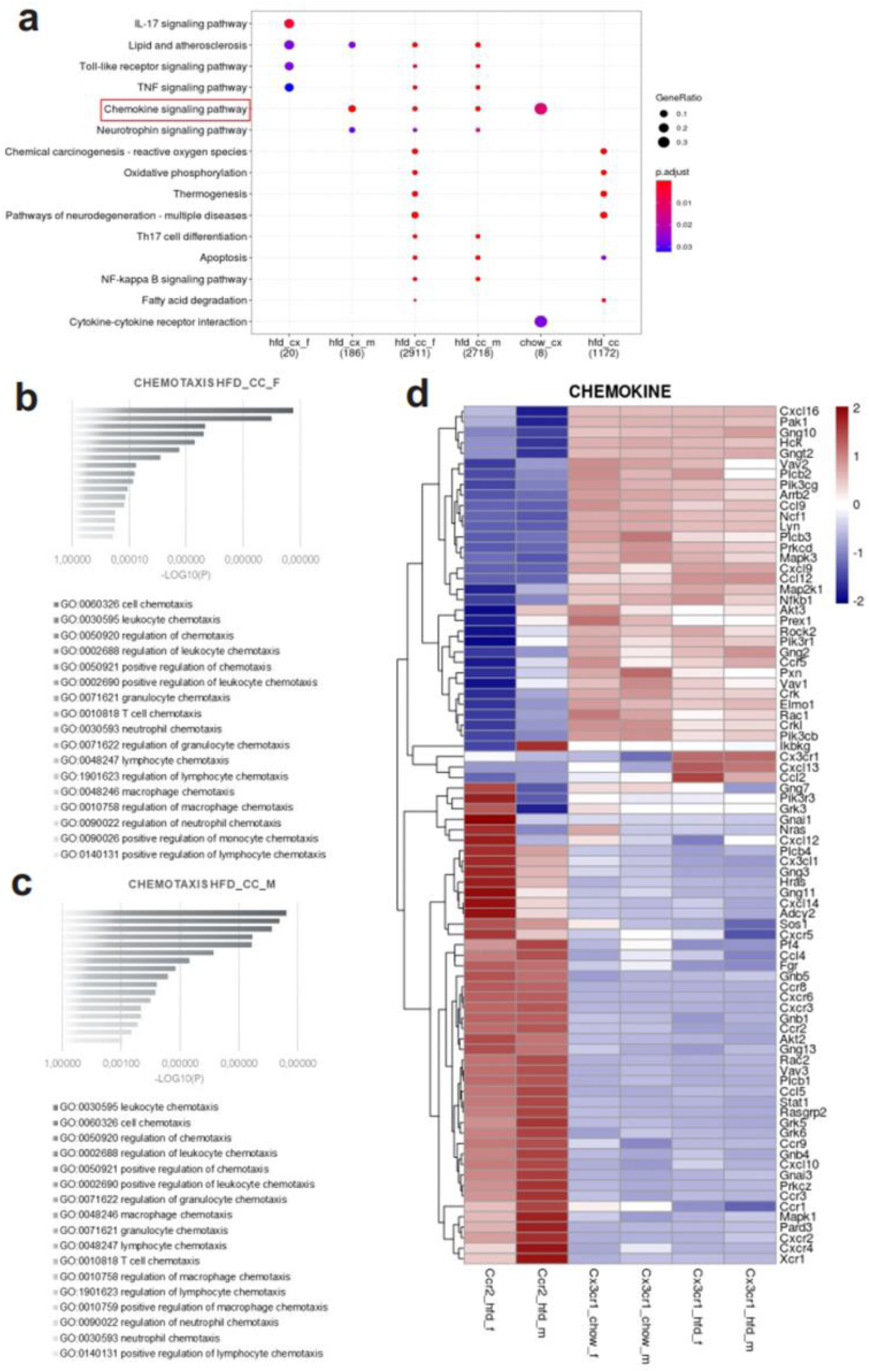
Various differential gene expression (DGE) found in CCR2+ infiltrating cells from the hypothalamus of HFD-fed mice belong to chemotaxis pathways. a) KEGG enrichment analysis shows the distribution of DGEs in distinct metabolic pathways. b-c) Ontology analysis for DGEs related to chemotaxis from CCR2+ cells sorted from the hypothalamus of HFD-fed female and male mice, respectively. d) Heatmap of up and downregulated DEGs related to chemotaxis when comparing CX3CR1+ resident and CCR2+ infiltrating cells from HFD-fed male and female mice.

**Table 1.**
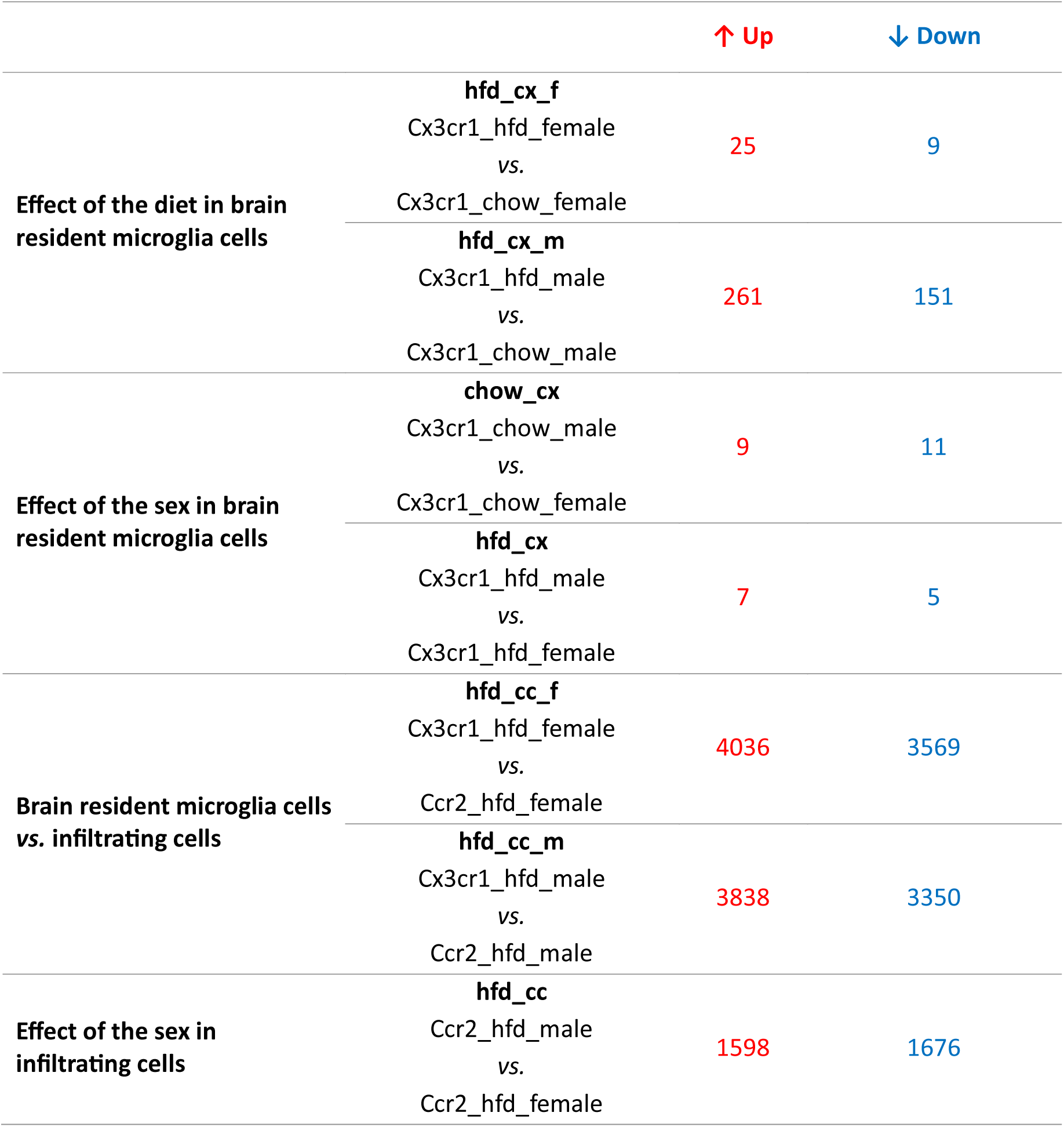
Number of DEGs for each comparison.

|  |  | ↑ Up | ↓ Down |
| --- | --- | --- | --- |
| **Effect of the diet in brain resident microglia cells** | **hfd_cx_f** |  |  |
|  | Cx3cr1_hfd_female | 25 | 9 |
|  | vs. |  |  |
|  | Cx3cr1_chow_female |  |  |
|  | **hfd_cx_m** |  |  |
|  | Cx3cr1_hfd_male | 261 | 151 |
|  | vs. |  |  |
|  | Cx3cr1_chow_male |  |  |
| **Effect of the sex in brain resident microglia cells** | **chow_cx** |  |  |
|  | Cx3cr1_chow_male | 9 | 11 |
|  | vs. |  |  |
|  | Cx3cr1_chow_female |  |  |
|  | **hfd_cx** |  |  |
|  | Cx3cr1_hfd_male | 7 | 5 |
|  | vs. |  |  |
|  | Cx3cr1_hfd_female |  |  |
| **Brain resident microglia cells vs. infiltrating cells** | **hfd_cc_f** |  |  |
|  | Cx3cr1_hfd_female | 4036 | 3569 |
|  | vs. |  |  |
|  | Ccr2_hfd_female |  |  |
|  | **hfd_cc_m** |  |  |
|  | Cx3cr1_hfd_male | 3838 | 3350 |
|  | vs. |  |  |
|  | Ccr2_hfd_male |  |  |
| **Effect of the sex in infiltrating cells** | **hfd_cc** |  |  |
|  | Ccr2_hfd_male | 1598 | 1676 |
|  | vs. |  |  |
|  | Ccr2_hfd_female |  |  |

### The impact of ovariectomy on diet-induced hypothalamic inflammation

Because sex exerted a major effect on the transcriptional signature of CCR2 cells, we evaluated ovariectomized mice fed on HFD, as depicted in the experimental design panel (Suppl. Fig. 1a). The ovariectomized mice presented increased body mass gain (Suppl. Fig. 1b) accompanied by increased adiposity (Suppl. Fig. 1e). This was accompanied by no change in food intake (Suppl. Fig. 1c), and blood glucose (Suppl. Fig. 1d). Regarding the hypothalamic expression of chemokines, there was only a reduction of Cxcl16 (Suppl. Fig. 1f), and no change in the expression of chemokine receptors (Suppl. Fig. 1g). Regarding the hypothalamic expression of neuropeptides involved in energy balance, there was an increase in Pomc and a reduction in Agrp (Suppl. Fig. 1h). There were no changes in the hypothalamic transcripts of inflammatory markers (Suppl. Fig. 1i). The blood levels of estradiol were determined (Suppl. Fig. 1j).

### Cxcr3 is highly expressed in recruited peripheral immune cells

As we were particularly interested in identifying factors involved in the recruitment of CCR2 to the hypothalamus, we evaluated cytokine receptors with high expression in CCR2 and low expression in CX3CR1. As depicted in Fig. 5, Ccr3 (Fig. 5a), Ccr7 (Fig. 5b), Ccr8 (Fig. 5c), Cxcr2 (Fig. 5d), Cxcr3 (Fig. 5e), Cxcr4 (Fig. 5f), Cxcr5 (Fig. 5g), Cxcr6 (Fig. 5h) and Cxcr7 (Fig. 5i) were all expressed in CCR2 cells and virtually absent from CX3CR1 cells. Cxcr3 (Fig. 5e) and Cxcr6 (Fig. 5h) presented the highest expressions, and therefore, we performed a search for previous studies looking at either of these chemokine receptors in the context of DIO hypothalamic inflammation. Using the terms, hypothalamus, hypothalamic, obesity, inflammation, Cxcr3, and Cxcr6, we could find no prior publications. As Cxcr3 is involved in interferon-gamma (IFN-γ) induction (Cole et al., 1998), and IFN-γ has been shown to be expressed in the context of DIO hypothalamic inflammation (De Souza et al., 2005), we looked into the IFN-γ-related pathways regulated in recruited immune cells (Fig. 6). First, we asked if the canonical ligands for CXCR3, C*XCL9*, CXCL10, and CXCL11, were expressed in either CX3CR1 or CCR2 cells. C*XCL11* was not detected in either cell type (not shown). CXCL9 (Fig. 6a) was expressed in CX3CR1 cells, only, whereas CXCL10 (Fig. 6b) was expressed in both CX3CR1 and CCR2 cells. In addition, in both female and male mice, Ifng was expressed in CCR2, but not in CX3CR1 cells (Fig. 6c). Furthermore, IFN-γ pathways were shown to be modulated in both female (Fig. 6d) and male (Fig. 6e) CCR2 cells. Thus, we elected CXCR3 as a target for further intervention.

**Figure 5.**
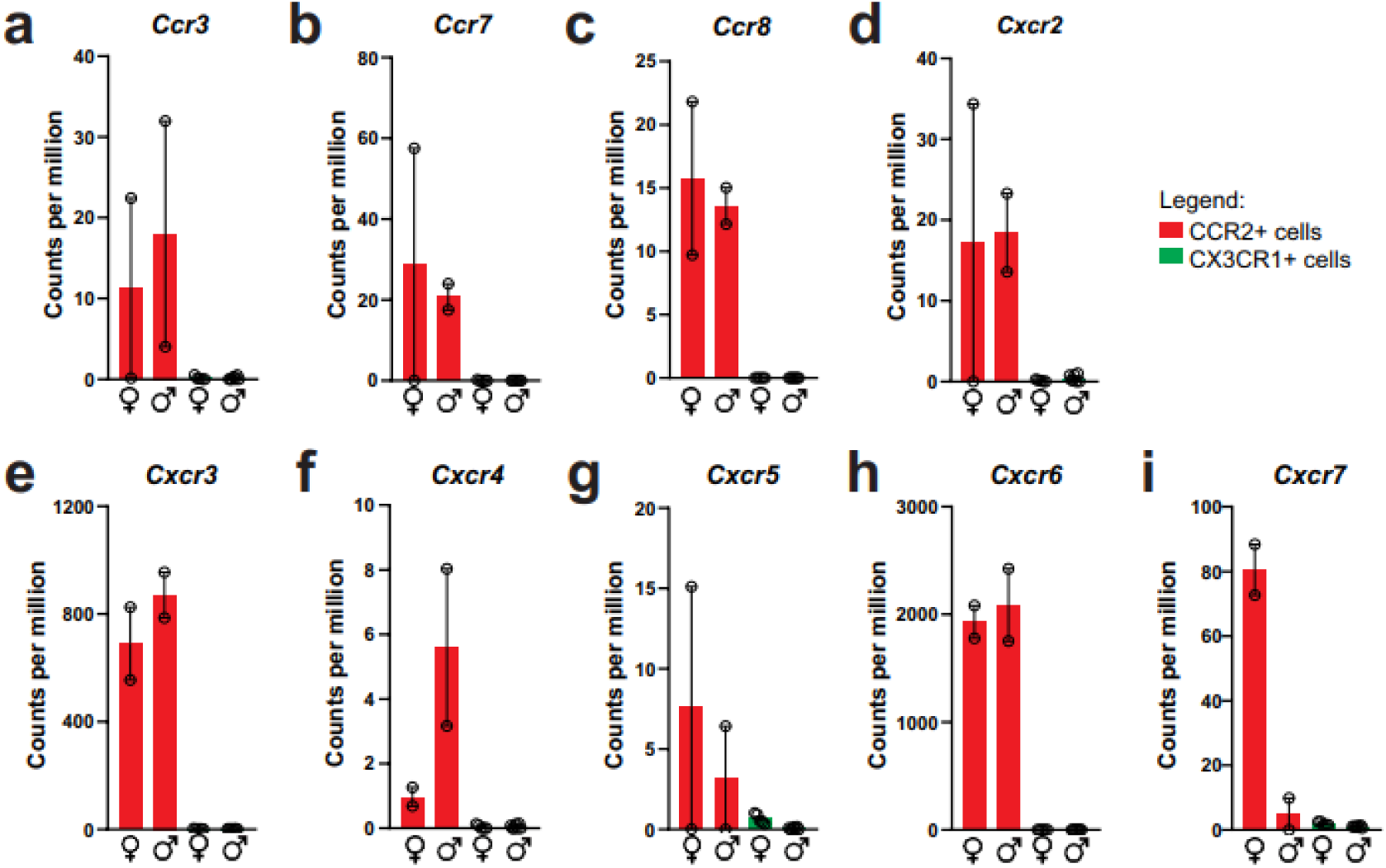
CCR2+ infiltrating cells from the hypothalamus of HFD-fed mice express a broad of chemokine receptors. a-i) Chemokine receptors gene expression in the transcriptome of CX3CR1+ and CCR2+ cells sorted from the hypothalamus of HFD-fed mice.

**Figure 6.**
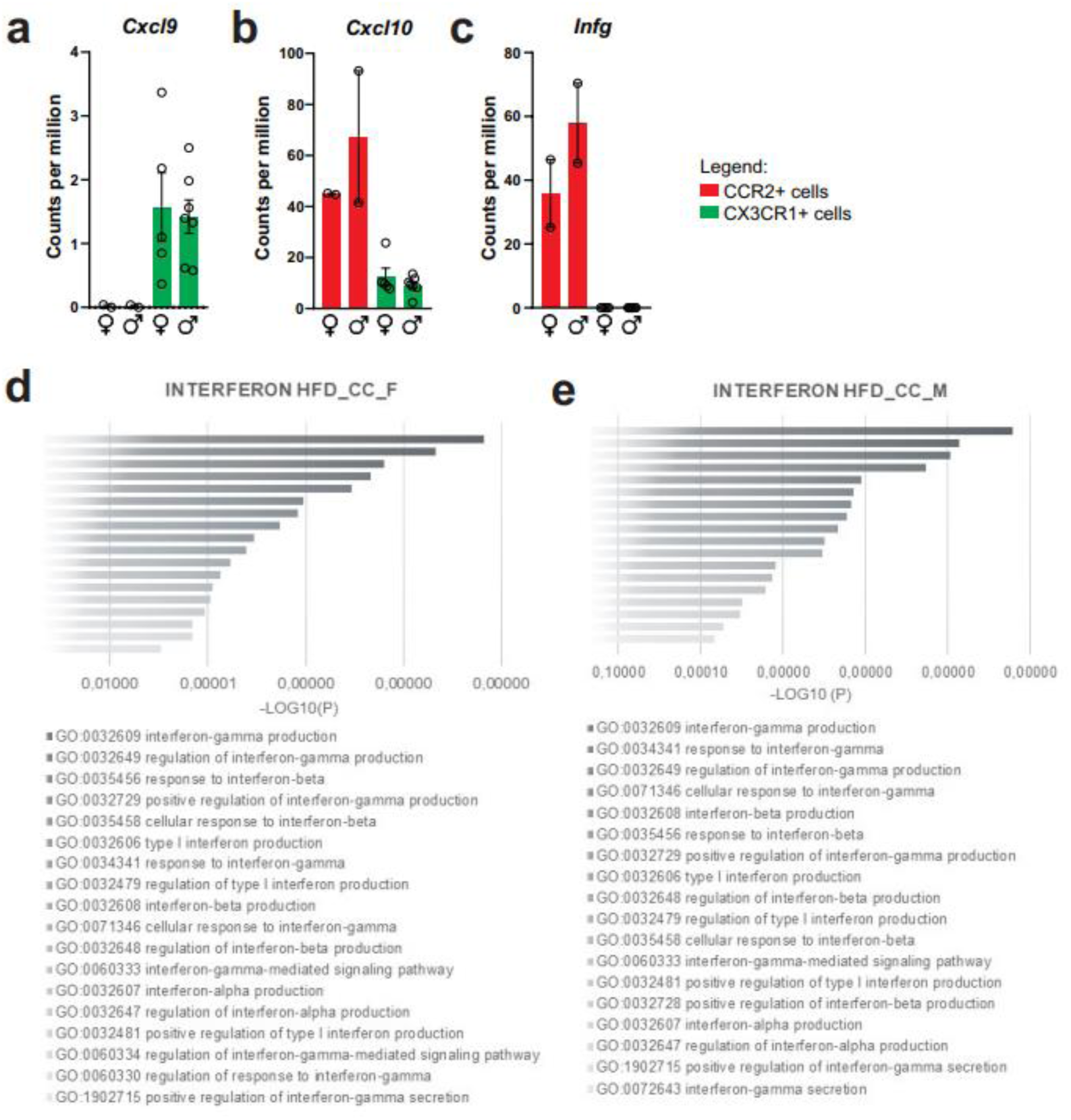
CXCL10/interferon γ-induced protein 10 kDa (IP-10) is highly expressed in CCR2+ infiltrating cells from the hypothalamus of HFD-fed mice. a-c) Cxcl9, Cxcl10 and Ifng gene expression in the transcriptome of CX3CR1+ and CCR2+ cells sorted from the hypothalamus of HFD-fed mice. d-e) Ontology analysis for DGEs related to interferon signaling pathways from CCR2+ cells sorted from the hypothalamus of HFD-fed female and male mice, respectively.

### The immunoneutralization of hypothalamic CXCL10 leads to increased body mass gain in female mice

As an attempt to interfere with CXCR3 actions in CCR2 cells, we targeted one of its ligands, CXCL10. As depicted in Fig. 7a, mice were submitted to two intracerebroventricular (icv) injections of an anti-CXCL10 antibody aimed at immunoneutralizing the target protein in the hypothalamus. As a result of the immunoneutralization of CXCL10, there were smaller numbers of CCR2-positive cells in the hypothalamus of both female and male mice fed on a HFD (Fig. 7b). However, there were no major changes in the numbers of CXCR3-expressing cells in the hypothalamus of either female (Fig. 7c) or male (Fig. 7d) mice fed on HFD. This was accompanied by no changes in the transcript levels of Cxcr3 and several other chemokine-related transcripts (Fig. 7c-7d), except for a trend to decrease Cxcl11 and an increase of Cxcr4 in females (Fig. 7c); and a decrease of Cxcl10 and a trend to increase Cx3cl1 in males (Fig. 7d). Nevertheless, the immunoneutralization of hypothalamic CXCL10 (Fig. 8 and Fig. 9) resulted in increased body mass gain (Fig. 8a-8b), a trend to reduce blood triglycerides (Fig. 8f), reduced blood cholesterol (Fig. 8g), reduced expression of Agrp transcript in the hypothalamus (Fig. 8k), a trend to reduce blood insulin (Fig. 8n), trends to reduce Il1b and Il6 transcripts in the hypothalamus (Fig. 8o), and a trend to reduce respiratory quotient (Fig. 8s) during the dark cycle in female mice. Conversely, in male mice (Fig. 9), the inhibition of hypothalamic CXCL10 had only a minor effect, leading to a trend to reduce hypothalamic Npy (Fig. 9k), a trend to reduce hypothalamic Il1b, and a trend to increase hypothalamic Il6 (Fig. 9o).

**Figure 7.**
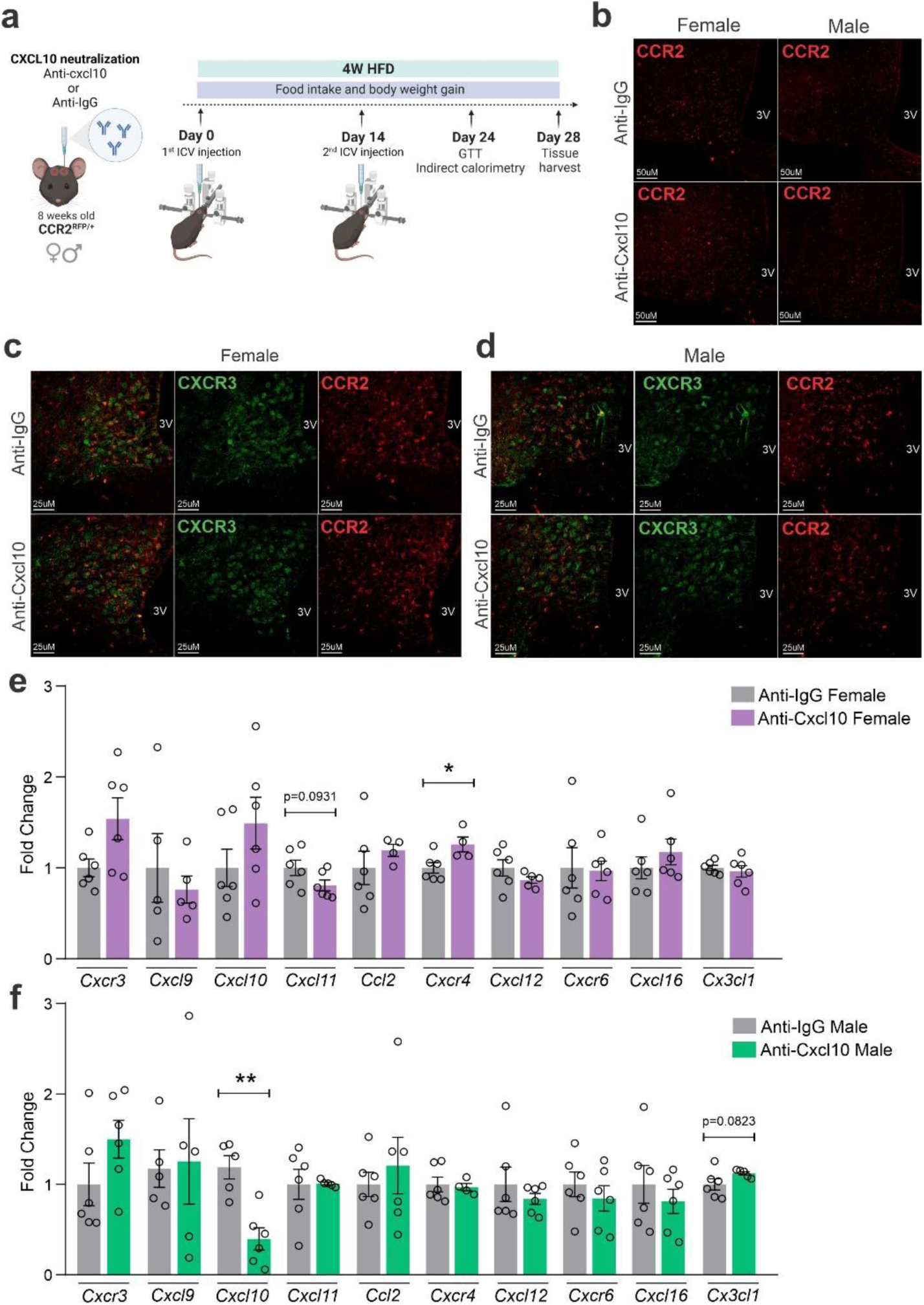
CXCL10 neutralization has a mild impact on reducing CCR2+ and CXCR3+ cell chemotaxis towards the hypothalamus of HFD-fed mice. a) Schematic representation of the experimental protocol for CXCL10 central neutralization. B) Coronal brain sections from 4 weeks HFD-fed CCR2^RFP^ mice showing the CCR2+ cells distribution in the hypothalamic parenchyma upon CXCL10 central neutralization. 3V = Third ventricle, Scale Bars = 50 μm. c) Coronal brain sections from 4 weeks HFD-fed female CCR2^RFP/+^ mice immunostained for CXCR3 upon CXCL10 central neutralization. D) Coronal brain sections from 4 weeks HFD-fed male CCR2^RFP/+^ mice immunostained for CXCR3 upon CXCL10 central neutralization. 3V = Third ventricle, Scale Bars = 25 μm. e-f) Hypothalamic Mrna levels of several chemokine receptors and chemokines in HFD-fed female (light gray and purple bars) and male (light gray and green bars) CCR2^RFP/+^ mice upon CXCL10 central neutralization. For qualitative confocal image analysis, we have used 3 samples per group. For RT-Qpcr of the hypothalamus, we have used 5-6 samples per group. Two-tailed Mann-Whitney tests were used for statistical analyses. *p<0.05 and **p<0.01 in comparison with respective Anti-IgG treated groups.

**Figure 8.**
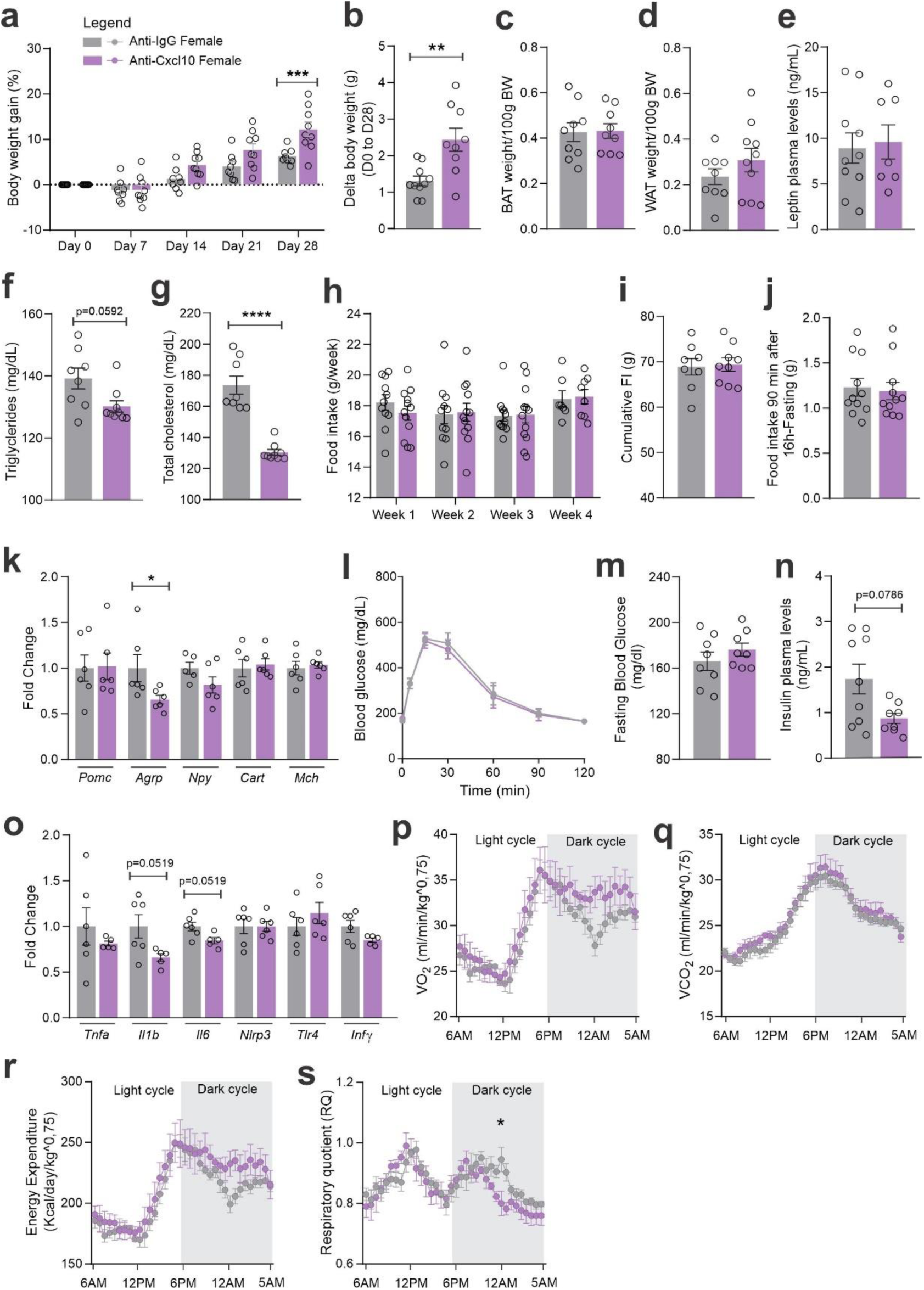
CXCL10 central neutralization in HFD-fed female mice. a) Percentual of body weight gain from Day 0 to Day 28 of the experimental protocol. b) Delta body weight during the experimental period. c) Brown adipose tissue weight and d) White adipose tissue (retroperitoneal depot) weight at Day 28. e) Leptin, f) Triglycerides, and g) Total cholesterol plasma levels at Day 28. h) Weekly food intake measurement during the experimental period. i) Cumulative food intake during the experimental period. j) 90-minute food intake measurement after 16 hours of fasting. k) Hypothalamic mRNA levels of neuropeptides involved in food intake control. l) Intraperitoneal glucose tolerance test on Day 24. m) 6h-fasting blood glucose levels. n) Insulin plasma levels at Day 28. o) Hypothalamic mRNA levels of inflammatory genes. p) O_2_ consumption; q) CO_2_ production; r) Energy Expenditure and s) Respiratory Quotient at Day 24. Data were expressed as mean ± SEM of 8-10 mice per group (in two independent experiments). To perform qRT-PCR we have used 6 mice/group. To perform biochemical analysis in plasma we have used 8-10 mice/group. To perform ipGTT we have used 4 mice/group. To perform indirect calorimetry, we have used 4 mice/group. Two-way ANOVA followed by Sidak’s post-hoc test and Mann-Whitney test were used for statistical analyses. *p<0.05, **p<001, ***p<0.001, ****p<0.0001 in comparison with IgG treated group.

**Figure 9.**
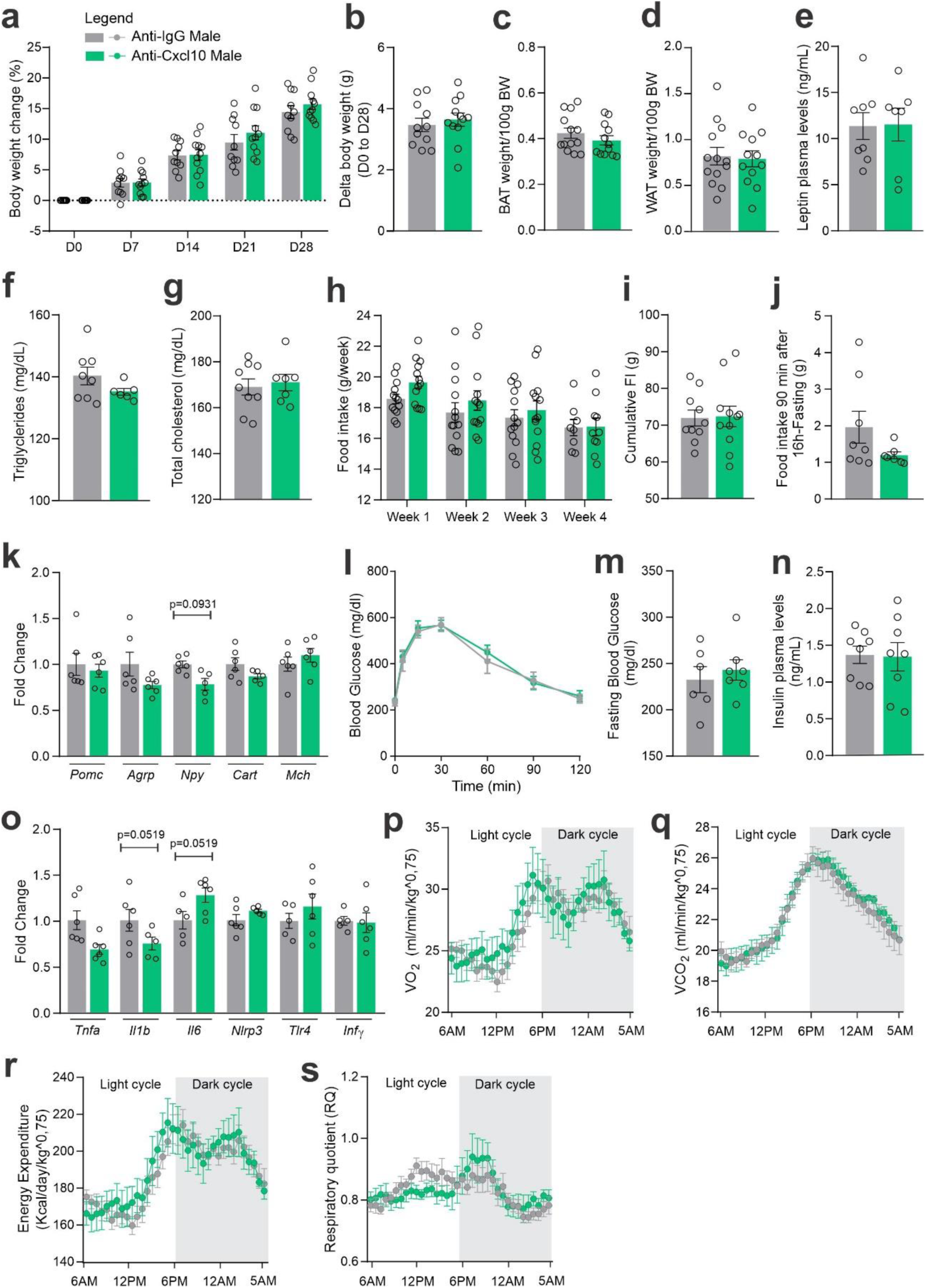
CXCL10 central neutralization in HFD-fed male mice. a) Percentual of body weight gain from Day 0 to Day 28 of the experimental protocol. b) Delta body weight during the experimental period. c) Brown adipose tissue weight and d) White adipose tissue (retroperitoneal depot) weight at Day 28. e) Leptin, f) Triglycerides and g) Total cholesterol plasma levels at Day 28. h) Weekly food intake measurement during the experimental period. i) Cumulative food intake during the experimental period. j) 90-minute food intake measurement after 16 hours of fasting. k) Hypothalamic mRNA levels of neuropeptides involved in food intake control. l) Intraperitoneal glucose tolerance test at Day 24. m) 6h- fasting blood glucose levels. n) Insulin plasma levels at Day 28. o) Hypothalamic mRNA levels of inflammatory genes. p) O_2_ consumption; q) CO_2_ production; r) Energy Expenditure and s) Respiratory Quotient at Day 24. Data were expressed as mean ± SEM of 8-10 mice per group (in two independent experiments). To perform qRT-PCR we have used 6 mice/group. To perform biochemical analysis in plasma we have used 8-10 mice/group. To perform ipGTT we have used 4 mice/group. To perform indirect calorimetry, we have used 4 mice/group. Two-way ANOVA followed by Sidak’s post-hoc test and Mann-Whitney test were used for statistical analyses. *p<0.05, **p<001, ***p<0.001, ****p<0.0001 in comparison with IgG treated group.

### The inhibition of CXCR3 worsens body mass gain and the metabolic phenotype of mice fed on a high-fat diet

CXCR3 was inhibited using a pharmacological antagonist, AMG487 ^24^ (Fig. 10a). We decided to perform the treatment systemically, instead of icv, because the purpose was to mitigate the migration of the cells expressing this receptor to the hypothalamus. Indeed, the intervention resulted in the reduction of CCR2 (Fig. 10b) and CXCR3 (Fig. 10c-10d) cells in the hypothalamus of both female and male mice fed on HFD. In addition, there was a reduction of Ccl2 and an increase of Cx3cl1 transcripts in the hypothalamus of female (Fig. 10e), and a reduction of Ccl2 transcripts in the hypothalamus of male (Fig. 10f) mice fed on HFD. The inhibition of CXCR3 had a major impact on metabolic phenotype; thus, in female mice fed a HFD, there was an increase in body mass gain (Fig. 11a-11b), a trend to increase brown adipose tissue mass (Fig. 11c), a trend to increase blood leptin (Fig. 11e), an increase in blood triglycerides (Fig. 11f), a trend to reduce hypothalamic Pomc (Fig. 11k), an increase in hypothalamic Npy (Fig. 11k), a worsen glucose tolerance (Fig. 11l), an increased fasting blood glucose (Fig. 11m), a reduced blood insulin (Fig. 11n), and reduction of Il6 and Tlr4 transcripts in the hypothalamus (Fig. 11o). In males, the inhibition of CXCR3 promoted an increased body mass gain (Fig. 12a-12b), increased white adipose tissue mass (Fig. 12d), increased blood leptin (Fig. 12e), increased hypothalamic Npy and Mch (Fig. 12k), and reduced hypothalamic Tnfa and Nlrp3 (Fig. 12o). Additionally, the inhibition of CXCR3 promoted changes in neither energy expenditure nor locomotor activity (Suppl. Fig 2a and 2b, for females and Suppl. Fig. 3a and 3b for males). In the hypothalamus of females there were no changes in the expression of transcripts encoding proteins involved in endoplasmic reticulum homeostasis (Suppl. Fig. 2c) and mitochondrial turnover (Suppl. Fig. 2d), whereas in males there was a reduction of Ddit3 (Suppl. Fig. 3c) and Mfn1 (Suppl. Fig. 3d). Moreover, in females the inhibition of CXCR3 promoted no changes in the liver expression of lipidogenic and gluconeogenic genes (Suppl. Fig. 2e), and no changes in the white adipose tissue expression of lipidogenic genes (Suppl. Fig. 2f). In the liver of males, there was a reduction in the expression of Fasn and an increase in the expression of G6pc3 (Suppl. Fig. 3e). As for the females, in males, there were no changes in the white adipose tissue expression of lipidogenic genes (Suppl. Fig. 3f).

**Figure 10.**
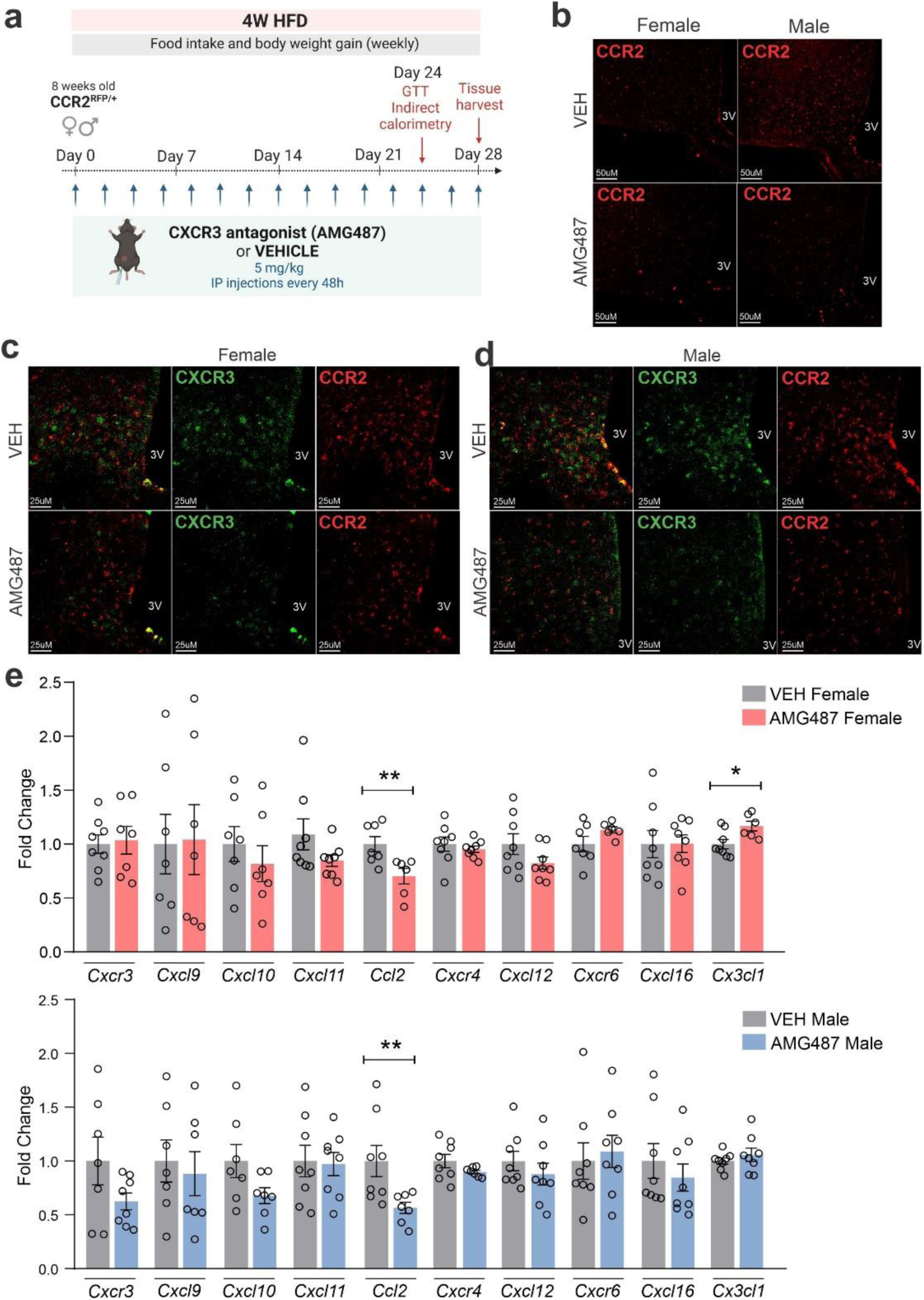
AMG487 treatment attenuates CCR2+ and CXCR3+ cell chemotaxis towards the hypothalamus of HFD-fed mice. a) Schematic representation of the experimental protocol for CXCR3 systemic blockage. b) Coronal brain sections from 4 weeks HFD-fed CCR2^RFP/+^ mice showing the CCR2+ cells distribution in the hypothalamic parenchyma upon AMG487 treatment. 3V = Third ventricle, Scale Bars = 50 μm. c) Coronal brain sections from 4 weeks HFD-fed female CCR2^RFP/+^ mice immunostained for CXCR3 upon AMG487 treatment. d) Coronal brain sections from 4 weeks HFD-fed male CCR2^RFP/+^ mice immunostained for CXCR3 upon AMG487 treatment. 3V = Third ventricle, Scale Bars = 25 μm. e-f) Hypothalamic mRNA levels of several chemokine receptors and chemokines in HFD-fed female (light gray and pink bars) and male (light gray and blue bars) CCR2^RFP/+^ mice upon AMG487 treatment. For qualitative confocal image analysis, we have used 3 samples per group. For RT-qPCR of the hypothalamus, we have used 7-8 samples per group. Two-tailed Mann-Whitney tests were used for statistical analyses. *p<0.05 and **p<0.01 in comparison with respective VEH-treated groups.

**Figure 11.**
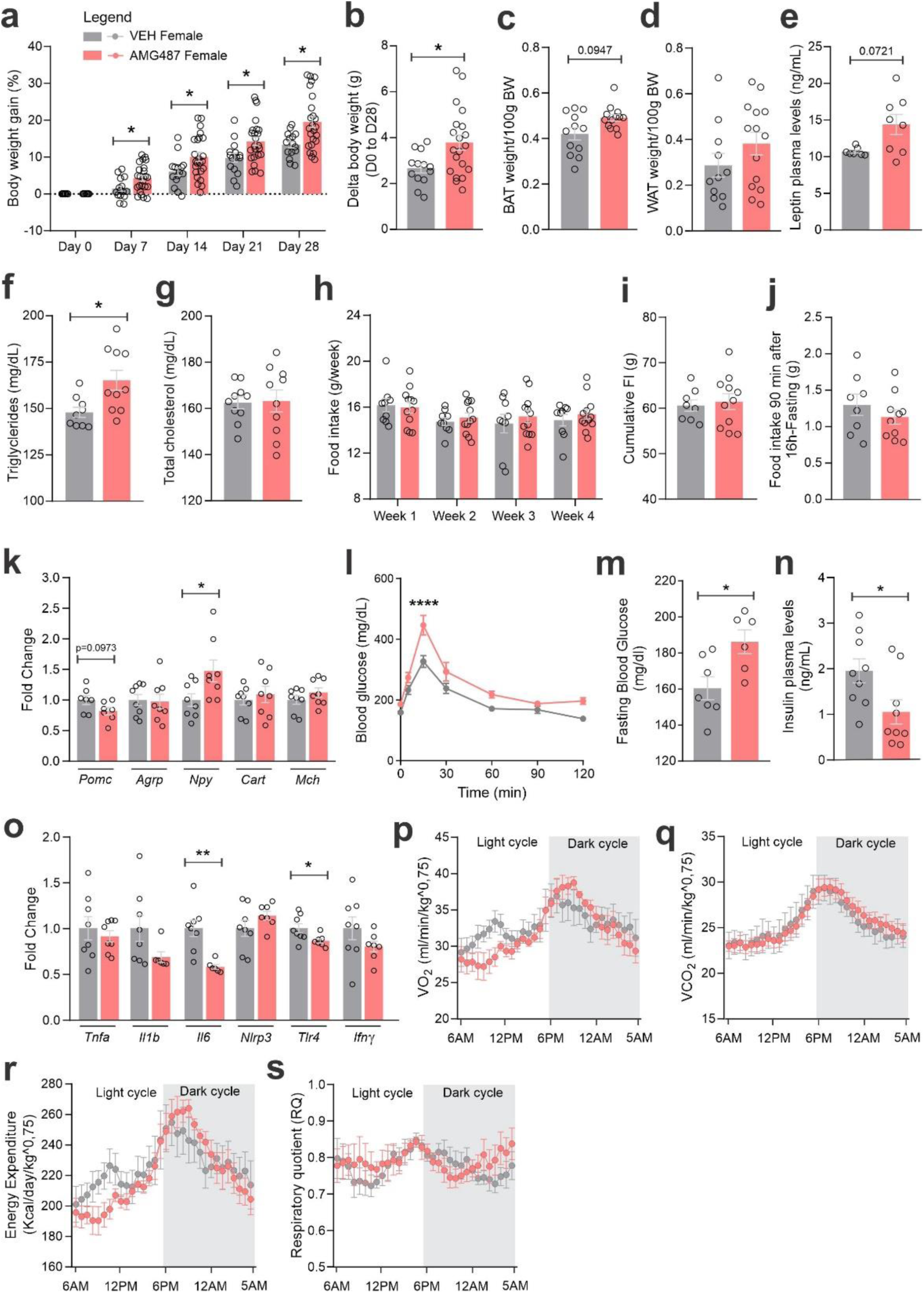
CXCR3 systemic blockage in HFD-fed female mice. a) Percentual of body weight gain from Day 0 to Day 28 of the experimental protocol. B) Delta body weight during the experimental period. C) Brown adipose tissue weight and d) White adipose tissue (retroperitoneal depot) weight at Day 28. E) Leptin, f) Triglycerides and g) Total cholesterol plasma levels at Day 28. H) Weekly food intake measurement during experimental period. I) Cumulative food intake during the experimental period. J) 90 min food intake measurement after 16h-fasting. K) Hypothalamic mRNA levels of neuropeptides involved in food intake control. L) Intraperitoneal glucose tolerance test at Day 24. M) 6h-fasting blood glucose levels. N) Insulin plasma levels at Day 28. O) Hypothalamic mRNA levels of inflammatory genes. P) O2 consumption; q) CO2 production; r) Energy Expenditure and s) Respiratory Quotient at Day 24. Data were expressed as mean ± SEM of 14-16 mice per group (in four independent experiments). To perform qRT-PCR we have used 8 mice/group. To perform biochemical analysis in plasma we have used 8-10 mice/group. To perform ipGTT we have used 5 mice/group. To perform indirect calorimetry, we have used 4-5 mice/group. Two-way ANOVA followed by Sidak’s post-hoc test and Mann-Whitney test were used for statistical analyses. *p<0.05, **p<001, ***p<0.001, ****p<0.0001 in comparison with VEH treated group.

**Figure 12.**
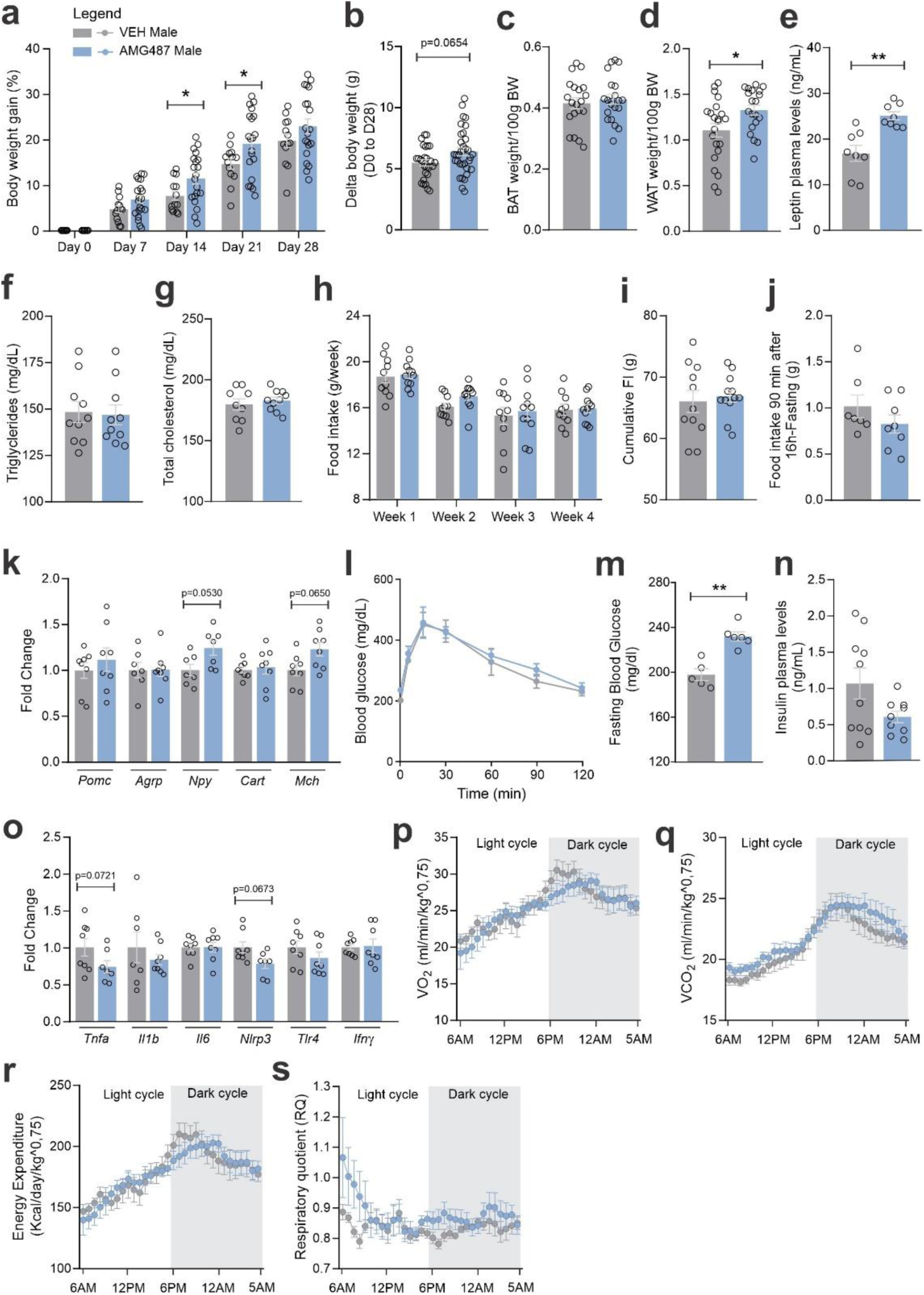
CXCR3 systemic blockage in HFD-fed male mice. a) Percentual of body weight gain from Day 0 to Day 28 of the experimental protocol. b) Delta body weight during the experimental period. c) Brown adipose tissue weight and d) White adipose tissue (retroperitoneal depot) weight at Day 28. e) Leptin, f) Triglycerides, and g) Total cholesterol plasma levels at Day 28. h) Weekly food intake measurement during the experimental period. i) Cumulative food intake during the experimental period. j) 90-minute food intake measurement after 16 hours of fasting. k) Hypothalamic mRNA levels of neuropeptides involved in food intake control. l) Intraperitoneal glucose tolerance test on Day 24. m) 6h-fasting blood glucose levels. n) Insulin plasma levels at Day 28. o) Hypothalamic mRNA levels of inflammatory genes. p) O2 consumption; q) CO2 production; r) Energy Expenditure and s) Respiratory Quotient at Day 24. Data were expressed as mean ± SEM of 14-16 mice per group (in four independent experiments). To perform qRT-PCR we have used 8 mice/group. To perform biochemical analysis in plasma we have used 8-10 mice/group. To perform ipGTT we have used 5 mice/group. To perform indirect calorimetry, we have used 4-5 mice/group. Two-way ANOVA followed by Sidak’s post-hoc test and Mann-Whitney test were used for statistical analyses. *p<0.05, **p<001, ***p<0.001, ****p<0.0001 in comparison with VEH treated group.

## Discussion

In this study, we elucidated the transcriptional landscapes of resident microglia and recruited immune cells in the hypothalamus of mice. We showed that resident microglia undergo minor transcriptional changes when mice are fed on HFD; however, there are vast differences when confronting resident CX3CR1+ microglia versus recruited CCR2+ immune cells transcriptomes. Moreover, there is a considerable degree of sexual dimorphism in the transcriptomes of recruited CCR2+ immune cells in mice fed a HFD. Upon exploration of the differences between resident microglia and recruited immune cells, we identified the chemokine receptor Cxcr3 as an interesting candidate for intervention as it was highly expressed in recruited cells. The inhibition of CXCR3 resulted in increased body mass gain, worsening of glucose intolerance, and increased expression of hypothalamic Npy. Thus, the study is the first to identify a subset of recruited immune cells that has a protective role against the deleterious outcomes of DIO, therefore establishing a new concept in obesity-associated hypothalamic inflammation.

Early studies in this field have shown that dietary fats, particularly long-chain saturated fatty acids, trigger an inflammatory response in the MBH that emerges a few hours after the introduction of a HFD and progresses to chronicity if the consumption of the HFD persists for long (De Souza et al., 2005) (Milanski et al., 2009) (Zhang et al., 2008). Microglia play an important role in this inflammatory response, and studies have shown that during the course of a prolonged consumption of HFD, there is the recruitment of bone marrow-derived cells (BMDCs) to compose a new hypothalamic microglia landscape (Thaler et al., 2012) (Morari et al., 2014) (Valdearcos et al., 2019). However, it was previously unknown what are the transcriptional signatures of hypothalamic resident microglia and recruited BMDCs in DIO.

To elucidate the transcriptional landscapes of resident microglia and recruited immune cells, we initially prepared a double reporter mouse for CX3CR1 and CCR2. CX3CR1 encodes for the fractalkine receptor and is highly expressed in resident microglia (Greter et al., 2015). The creation of CX3CR1 reporter mice was regarded as an important step toward the characterization of resident microglia, and several studies in the field employ this model (Goldmann et al., 2013) (Yona et al., 2013). Conversely, CCR2 is expressed only in bone marrow-derived cells (Greter et al., 2015) (Mildner et al., 2007). The quality of our model was proven good as markers of resident microglia were present in CX3CR1 cells, only, whereas markers of recruited peripheral immune cells were present in CCR2 cells, only. Moreover, as previously described, we could find no CCR2 cells in the hypothalamus of mice fed chow (Valdearcos et al., 2017) (Valdearcos et al., 2019).

The first important, and previously unknown finding emerged from the comparison of the transcriptomes of resident microglia in mice fed chow versus mice fed HFD. In females, there were only 34 transcripts undergoing significant changes between the two dietary conditions, whereas in males, the number was greater, 412, but still quite small considering the whole transcriptome of resident microglia (Young et al., 2021). In a study evaluating the single-cell transcriptomics of hypothalamic cells, the consumption of a HFD resulted in minor changes in the so-called macrophage-like cells (Campbell et al., 2017); however, the detailed transcriptional landscape of these cells was not explored in depth, so it is uncertain if it contained both resident microglia and recruited immune cells. In an experimental model of Alzheimeŕs disease, which evaluated male and female mice, there were hippocampal resident microglial transcriptional changes of the same magnitude as the one we found in the hypothalamus, affecting approximately 300 genes (Rivera-Escalera et al., 2019). In an experimental model of cerebral hemorrhage evaluating only males, the impact on microglia transcriptome was also small, affecting only 10% of the evaluated genes (Taylor et al., 2017). In addition, in a study evaluating transcriptional changes of resident microglia during aging (Li et al., 2023), there were also important differences between female and male mice; and, the magnitude of the transcriptional changes occurring during aging was about the same as we see in the dietary intervention. Thus, it seems that in different brain regions and under distinct interventions, the magnitude of resident microglia transcriptional changes is quite small; however, the number of studies evaluating this question is not expressive; thus, further studies are needed to provide a definitive view regarding the actual magnitude of plasticity of resident microglia. Despite the small number of transcripts undergoing changes in our model, greatest impact occurred in the expression of genes related to IL-17 signaling, lipid metabolism, TLR signaling, TNF signaling and chemokine signaling, which strongly supports the role of dietary lipids in the induction of an inflammatory response by the resident microglia, supporting previous studies in the field (Milanski et al., 2009) (Thaler et al., 2012) (Morari et al., 2014) (Valdearcos et al., 2019). Interestingly, IL17 signaling emerged as a major pathway modulated by DIO. In a recent study, we have shown that IL17 can act upon POMC neurons promoting a reduction in calorie intake (Nogueira et al., 2020). The current finding of a transcriptional modulation of IL17-related genes in resident microglia opens a new perspective in the understanding of how inflammatory signals modulate the function of the hypothalamus in obesity, which should be explored in the future.

Next, we confronted the transcriptomes of resident microglia and recruited immune cells, and in this case, there were huge differences, reaching over 7,000 transcripts. Per se, this finding reveals a striking difference between resident microglia and recruited immune cells in the hypothalamus of mice fed on HFD. Nevertheless, despite this is a new information regarding the hypothalamus, similar findings were reported in other brain regions submitted to distinct interventions, such as in an experimental model of glioma (Ochocka et al., 2021), in flavivirus infection (Spiteri et al., 2023), and in COVID-19 (Schwabenland et al., 2021) (Hartmann et al., 2024). The evaluation of the main pathways differentially expressed in the two cell subsets revealed major differences in lipids, toll-like receptor signaling, tumor necrosis factor signaling, chemokines, neurotrophins signaling, reactive oxygen species, thermogenesis, and pathways related to neurodegeneration. The impact of the consumption of a HFD on the regulation of lipid-related pathways, toll-like receptor, and tumor necrosis factor signaling has been widely explored in many previous studies, and interventions in these systems are known to mitigate the harmful effects of the diet (Milanski et al., 2009) (Thaler et al., 2012) (Morari et al., 2014) (Valdearcos et al., 2019). However, little has been done regarding the characterization of the mechanisms of chemotaxis that drive the recruitment of bone marrow-derived cells to the hypothalamus. Therefore, we looked with greater detail into the main chemokines and chemokine receptors differentially expressed in the two subsets of cells. We elected Cxcr3 because it was highly expressed in Ccr2 cells and presented low expression in Cx3cr1 cells. CXCR3 is a chemokine receptor that is involved in the recruitment of distinct types of bone marrow-derived monocytic cells, such as plasmacytoid monocytes (Cella et al., 1999), synovial tissue monocytes (Katschke et al., 2001), and dendritic cells (Padovan et al., 2002). In the brain, CXCR3-expressing cells have been implicated in Alzheimer’s disease and other age-dependent cognitive dysfunctions (Jorfi et al., 2023) (Schroer et al., 2023), multiple sclerosis (Bogers et al., 2023), epilepsy (Liang et al., 2023), and stoke (Cai et al., 2022). However, no previous study has evaluated CXCR3 in the hypothalamus in the context of obesity.

First, we asked if the known ligands for CXCR3, and Ifng, which is induced in response to the activation of CXCR3, were present in either subset of cells. We found Cxcl11 in neither cell subset; Cxcl9 was expressed in Cx3cl1 cells, only; Cxcl10 was expressed in both subsets of cells, with greater expression in Ccr2 cells; and Ifng was expressed in Ccr2 cells only. Moreover, there was a considerable engagement of IFN-γ-related pathways in Ccr2 cells of both female and male mice fed a HFD. Thus, we considered CXCR3 as a promising target for intervention.

Next, we intervened in one of the ligands for CXCR3, CXCL10. For that, we performed icv injections of an immunoneutralizing antibody, which resulted in smaller numbers of CCR2 cells in the hypothalamus. However, the intervention promoted minimal changes in the hypothalamic expression of transcripts encoding for several components of the chemotaxis machinery. Nevertheless, in females, the inhibition of CXCL10 resulted in increased body mass gain, reduction of hypothalamic Agrp, trends to reduce hypothalamic Il1b and Il6, and a trend to reduce blood insulin. In males, the phenotype was much milder, leading to minimal changes in hypothalamic Npy, Il1b, and Il6. Little is known about the involvement of CXCL10 in hypothalamic physiology and pathology. In a model of caloric restriction, there was an increase in the hypothalamic expression of Cxcl10 (Matthews et al., 2017), and this was regarded as a component of the mechanism of neuroprotection induced by caloric restriction. In addition, in a model of hypothalamic inflammation elicited by exogenous LPS, Cxcl10 emerged as one of the transcripts undergoing the greatest increase in the hypothalamic paraventricular nucleus (Reyes et al., 2003).

As CXCR3 can be engaged by distinct chemokines, we decided to use a broader intervention, pharmacologically inhibiting CXCR3. AMG487 is a chemical inhibitor of CXCR3 with an IC_50_ value of 8.0 nM (Johnson et al., 2007). Upon treatment with AMG487, there was a reduction of the migration of Ccr2 cells to the hypothalamus, which was accompanied by minimal changes in the expressions of chemokines and chemokine receptors. Under inhibition of CXCR3, both female and male mice presented increased body mass gain, which was accompanied by increased blood leptin, increased fasting glucose, increased hypothalamic Npy, and reduction in markers of hypothalamic inflammation. There were no changes in caloric intake, however, there were reductions in energy expenditure during some periods during the 24 hours of recording.

These findings reveal that, at least one subset of recruited immune cells, has a protective rather than a harmful role in DIO-associated hypothalamic inflammation. This is a completely new concept in the field because all previous studies evaluating hypothalamic microglia in DIO reported that, once active in response to dietary fats, either resident microglia or recruited immune cells exerted inflammatory actions that impacted negatively energy balance and glucose tolerance (Tapia-González et al., 2011) (Valdearcos et al., 2014) (Fernández-Arjona et al., 2022) (Morari et al., 2014). This concept has been recently explored in depth in a study that used elegant models to either activate or inactivate microglia (Valdearcos et al., 2017) and the results confirm that, whenever manipulating microglia using approaches that are not specific for a given subset of cells, the net result is worsening of the metabolic phenotype when microglia is activated and improvement of the metabolic phenotype when microglia is inactivated. Thus, we believe that the subset of recruited immune cells herein identified plays a regulatory role in hypothalamic inflammation.

We acknowledge that the study could be strengthened if, in addition to pharmacological and antibody-based interventions, we used gene-targeted approaches as well.

In conclusion, this study elucidated the transcriptional landscapes of hypothalamic resident microglia and recruited immune cells in DIO. In resident microglia, the consumption of a HFD resulted in small changes in transcript expression, whereas the confrontation of the transcriptional landscapes of resident microglia versus recruited immune cells revealed broad differences that encompass lipids, toll-like receptor signaling, tumor necrosis factor signaling, chemokines, neurotrophins signaling, reactive oxygen species, thermogenesis, and pathways related to neurodegeneration. In addition, the study revealed a considerable sexual dimorphism in the transcriptional landscape of both resident microglia and recruited immune cells. The study also identified a subset of recruited immune cells, expressing the chemokine receptor Cxcr3 that has a protective role against the harmful metabolic effects of the HFD; thus, providing a new concept in DIO-associated hypothalamic inflammation.

## Materials and methods

### Animal care and diets

All animal care and experimental procedures were conducted in accordance with the guidelines of the Brazilian College for Animal Experimentation and approved by the Institutional Animal Care and Use Committee (CEUA 5497-1/2020 and 6210-1/2023). Heterozygous CX3CR1^GFP/+^CCR2^RFP/+^ mice were generated by mating CX3CR1^GFP^ homozygous mice (JAX#005582) with CCR2^RFP/+ hetero^zygous mice (JAX#017586). Genotypes of these mice were identified by polymerase chain reaction (PCR). Mice were fed on standard chow diet (Nuvilab; 3.76 kcal/g; 12.6% energy from protein, 77.7% energy from carbohydrate, and 9.58% energy from fat) or high-fat diet (HFD) (5.28 kcal/g; 12.88% energy from protein, 27.1% energy from carbohydrate, and 60% energy from fat) according to the experimental protocols. Food and water were available ad libitum throughout the experimental periods, except for the protocols that required fasting. The room temperature was controlled (22-24 °C), and a light-dark cycle was maintained on a 12-hour on-off cycle.

### Flow cytometry

For the separation of CX3CR1^GFP+^ and CCR2^RFP+^ cells from the white adipose tissue of CX3CR1^GFP/+^CCR2^RFP/+^ mice we collected the retroperitoneal fat depot of one animal fed on a HFD for 4 weeks. It was minced and digested with type VIII collagenase (0.5 mg/mL, Sigma-Aldrich) in PBS for 20 min at 37°C with shaking. After digestion, the suspension was filtered using a 100 μm cell filter. For isolation of the same cells from the hypothalamus, samples of five CX3CR1^GFP/+^CCR2^RFP/+^ mice fed on a HFD for 4 weeks were pooled together and gently pressed through a cell strainer (100 μm). The cell solution was subjected to a Percoll gradient (70/40%) for monocyte purification. Samples were acquired on a BDFacs Symphony instrument (BD Biosciences, USA) and then analyzed using FlowJo software.

### Cell sorting

For cell sorting of CX3CR1^GFP+^ and CCR2^RFP+^ cells from the hypothalamus we employed CX3CR1^GFP/+^ mice fed on chow diet and CX3CR1^GFP/+^CCR2^RFP/+^ mice fed on HFD for 4 weeks. Harvested hypothalami of 20-30 male or 20-30 female mice were pooled together for each sample and gently pressed through a cell strainer (100 μm). The cell solution was subjected to a Percoll gradient (70/40%) for monocyte purification. The sorting was conducted on a BDFacs Melody instrument (BD Biosciences, USA).

### RNA-sequencing (RNA-Seq) and analysis

Cell-sorted CX3CR1 GFP+ and CCR2 RFP+ cell samples from hypothalamus were lysed for RNA extraction using the RNAqueous Micro kit (Invitrogen). RNA integrity was analyzed on a Bioanalyzer RNA Pico 6000 chip at the Core Facility for Scientific Research – University of São Paulo (CEFAP-USP). Low input RNA-Seq library preparation (Takara SMART-Seq v4) and sequencing by Illumina NovaSeq S2 PE150 Sequencing Lane (40M read pairs/sample avg) were performed by Maryland Genomics (Institute for Genome Sciences - IGS, University of Maryland School of Medicine – Baltimore, USA). Illumina sequencing adapters and low-quality reads were removed with Trimmomatic. Trimmed reads were aligned to the mouse reference genome (GRCm39) by STAR. Aligned reads were mapped to features using HTSeq, and differential expression analyses were performed using the DESeq2 package. Genes having less than 3 CPM were excluded before statistical analysis, and differentially expressed genes (DEGs) were selected using as cutoffs the adjusted p-value < 0.05. Heatmaps were performed using pheatmap and a list of DEGs was passed to enrichR and cluster Profiler for enrichment analyses.

### CXCL10 immunoneutralization

For central neutralization of CXCL10, 8-wk CCR2^RFP/+^ male and female mice underwent a stereotaxic surgery for ICV injections of anti-CXCL10 Monoclonal Antibody (2 ul, Cat# MA5-23774, Thermo Fischer). The control groups were ICV injected with Mouse IgG2a Isotype Control (2 ul, Cat#02-6200, Thermo Fischer). Two distinct ICV injections were performed on Day 0 and Day 14, respectively, of the experimental protocol. For that, mice were anesthetized with ketamine (100 mg/kg) and xylazine (10 mg/kg) and submitted to stereotaxic surgery (Ultra Precise–model 963, Kopf). ICV coordinates were [antero-posterior/lateral/depth to bregma]: −0.46/−1.0/−2.3 mm. Immediately after the first surgery, at Day 0, mice began to be fed on HFD for 4 weeks. From Day 0 to Day 28 food intake and body weight were evaluated weekly.

### CXCR3 antagonism

For systemic blockage of CXCR3, 8-wk CCR2^RFP/+^ male and female mice underwent a treatment with AMG487 (Tocris Bioscience, Bristol, UK), an active and selective CXC chemokine receptor 3 (CXCR3) antagonist. The in vivo formulation of AMG487 was prepared in 20% hydroxypropyl-β-cyclodextrin (Sigma, St. Louis, MO). A 50% hydroxypropyl-β-cyclodextrin (Sigma, St. Louis, MO) solution was prepared and AMG487 was added to this solution, it was incubated in a sonicating water bath for 2 hours with occasional vortexing.

Next, distilled water was added to give the appropriate final concentration of AMG487 in 20% of hydroxypropyl-β-cyclodextrin. This solution at 20% served as the vehicle. Mice were treated with AMG487 or vehicle (VEH group) intraperitoneally at 5 mg/kg every 48h throughout four weeks. During this period, mice were fed on HFD, and food intake and body weight were evaluated weekly.

### Ovariectomy procedure and estradiol replacement

Female C57BL/6J mice were anesthetized with ketamine (100 mg/kg) and xylazine (10 mg/kg). The ventral abdominal area was shaved and sterilized using an iodine solution. A small midline incision was made, and the ovaries were carefully located and excised bilaterally. The incision was then sutured using a suture thread. The Sham group, which underwent the same procedure except for ovary excision, was used as the control. Half of the ovariectomized mice also received estradiol replacement therapy. 17β-estradiol pellets (0.05 mg/pellet, 60-day sustained release; Innovative Research of America, Inc., USA) were implanted subcutaneously beneath the dorsal surface of the neck. Tramadol hydrochloride (5 mg/kg, intraperitoneally) was administered immediately post-surgery, as well as 24 hours and 36 hours after surgery to manage pain. Mice were monitored during a 7-day recovery period. From Day 7 to Day 35, all groups were fed a high-fat diet (HFD), and food intake and body weight were evaluated weekly. On the 25th day of the protocol, mice were fasted for 6 hours. Before euthanasia, we measured fasting glycemia. Afterward, we harvested the hypothalamus and the retroperitoneal white adipose tissue (WAT). The WAT was weighed for adiposity measurement, and the hypothalamus was used for qPCR analysis of chemokines, chemokine receptors, neuropeptides, and some inflammatory markers.

### Glucose tolerance test

On the 24th day of CXCR3 blockage and CXCL10 neutralization experimental protocols, mice were fasted for 6 hours, and blood glucose was measured via tail bleed at baseline and 15, 30, 60, 90, and 120 min after an intraperitoneal injection of glucose (2.0 g/kg).

### Indirect calorimetry and locomotor activity

The oxygen consumption (VO_2_), carbon dioxide production (VCO_2_), energy expenditure, and respiratory quotient (RQ) were measured using an indirect open-circuit calorimeter (Oxylet M3 system; PanLab/Harvard Apparatus, MA, USA).

Spontaneous locomotor activity was measured using a Panlab Infrared (IR) Actimeter, which consists of a two-dimensional (X and Y axes) square frame, a frame support, and a control unit. Each frame is equipped with 16 x 16 infrared beams for optimal subject detection (PanLab/Harvard Apparatus, MA, USA). For each mouse, we calculated the mean of total movements per hour over 24 hours. Mice were allowed to adapt for 12 h before data were recorded for 24 h (light and dark cycles).

### Immunofluorescence

On the day 28th of CXCR3 blockage and CXCL10 neutralization experimental protocols, male and female mice were perfused with 0,9% saline followed by 4% formaldehyde by cardiac cannulation. Brains were extracted and incubated in 4% formaldehyde overnight at 4 °C for extended fixation. The brains were then incubated in 30% sucrose at 4 °C for 48h. A series of 20 μm-thick frozen sections (4 series equally) were prepared using a cryostat and stored in an anti-freezing solution. For the free-floating immunostaining, slices were washed with 0.1 M phosphate-buffered saline (PBS) (3 times, 5 min each) and blocked with 0.2% Triton X-100 and 5% donkey serum in 0.1 M PBS for 2 h at room temperature. Slices were incubated overnight at 4 °C with Anti-Cxcr3 (1:200, Cat# NB100-56404, Novus Biologicals) or Anti-Sialoadhesin/CD169 (1:200, ab18619, Abcam) in a blocking solution. After washing with 0.1 M phosphate-buffered saline (PBS) (3 times, 5 min each), sections were incubated with fluorophore-labeled secondary antibody (donkey anti-rabbit Alexa Fluor 405, 1:500, Cat# A48258, Invitrogen or goat anti-mouse Alexa Fluor 405, 1:500, Cat# A31553, Invitrogen) in a blocking solution for 2 h at room temperature. After washing again with 0.1 M phosphate-buffered saline (PBS) (3 times, 5 min each), brain slices were mounted onto slides with ProLong Diamond antifade mountant (Cat# P36930, Thermo Fischer). Sections were visualized with a Zeiss LSM780, confocal microscope (Carl Zeiss AG, Germany) at the National Institute of Photonics Applied to Cell Biology (INFABIC) at the University of Campinas.

### Quantitative reverse transcription-polymerase chain reaction (qRT-PCR)

Total RNA was extracted using a TRIzol reagent (Thermo Fisher Scientific) and synthesized cDNA with a High-Capacity cDNA Reverse Transcription Kit (HighCapacity cDNA Reverse Transcription Kit, Life Technologies). Real-time PCR reactions were run using the TaqMan system (Thermo Fisher Scientific). Primers used were Cxcr3 (Mm99999054_s1); Cxcl9 (Mm00434946_m1), Cxcl10 (Mm00445235_m1); Cxcl11 (Mm00444662_m1); Ccl2 (Mm00441242_m1); Cxcr4 (Mm01996749_s1); Cxcl12 (Mm00445553_m1), Cxcr6(Mm02620517_s1), Cxcl16 (Mm00469712_m1), Cx3cl1(Mm00436454_m1), Pomc (Mm00435874_m1), Agrp (Mm00475829_g1); Npy (Mm00445771_m1); Cartpt (Mm04210469_m1), Pmch (Mm01242886_g1), Tnfa (Mm00443258_m1), Il1b (Mm00434228_m1), Il6 (Mm00446190_m1), Nlrp3 (Mm00840904_m1), Tlr4 (Mm00445273_m1), Infg (Mm01168134_m1). Eif2a (Mm01289723_m1), Atf6 (Mm01295319_m1), Ddit3 (Mm01135937_g1), Immp2l (Mm00474144_m1), Mfn1 (Mm00612599_m1), Opa1 (Mm01349707_g1), Htra2 (Mm00444846_g1), Ppargc1a (Mm01208835_m1), FasN (Mm00662319_m1), Scd1 (Mm00772290_m1), Scd2 (Mm01208542_m1), Pck1 (Mm01247058_m1), G6pc3 (Mm00616234_m1), G6pc (Mm04207416_m1), Pparg (Mm00440940_m1), Prdm16 (Mm00712556_m1), Ucp1 (Mm01244861_m1). GAPDH (Mm99999915_g1) was employed as the reference gene for all tissues, except for white adipose tissue, where ACTB (Mm02619580_g1) was employed as the reference gene. Gene expression was obtained using QuantStudio 6 (Thermo Fischer Scientific).

### Hormonal and biochemical determinations

Serum insulin, and leptin were measured by an enzyme-linked immunosorbent assay (ELISA) kit (#EZRMI-13K and #EZML-82K; Millipore; E-EL—0150; Elabscience). Serum triglyceride levels and total cholesterol were measured using a commercial colorimetric assay kit (LaborLab®, Guarulhos - SP, Brazil) following the manufacturer’s instructions.

### Statistical analysis

Data are presented as means ± standard error of the mean (SEM). The statistical analyses were carried out using a non-parametric Mann-Whitney test and one- or two-way analysis of variance (ANOVA) when appropriate. Post hoc comparisons were performed using Sidak’s test. Statistical significances were analyzed using Prism 8.0 software (GraphPad Software, La Jolla, CA). A p-value ≤0.05 was considered statistically significant. In the experiments aimed at measuring energy expenditure, data was always corrected for body mass.

## Declaration of Interests/Relevant conflicts of interests/Financial disclosures

*All* authors declare no competing financial or other interests regarding this study.

## Authors’ contributions

NFM, EPA, and LAV designed and planned the study; NFM, AMZ, DCS, and JFC performed most of the experiments; NFM and CFA performed FACS and cytometry analysis; GFL conducted bioinformatic analysis; NOSC provided animal models; PMMM-V supervised the work of CFA and provided equipment and methods for FACS and cytometry; NFM, EPA, and LAV discussed the data. NFM and LAV wrote the manuscript and prepared figures; EPA supervised NFM during her doctorate program. All authors read and approved the final manuscript.

## Funding

This research was funded by The Sao Paulo Research Foundation (FAPESP): 2013/07607-8; 2017/22511-8; and 2021/00443-6.

## Data Availability

Data will be available upon request to the corresponding author, L.A.V. The RNA sequencing data for this study were submitted to the NCBI Sequence Read Archive (SRA) under BioProject accession number PRJNA1155598.

## Acknowledgments

We are grateful to Erika Roman, Joseane Morari, Marcio Cruz, and Gerson Ferraz for laboratory management.

## SUPPLEMENTARY FIGURES

**Supplementary Figure 1.**
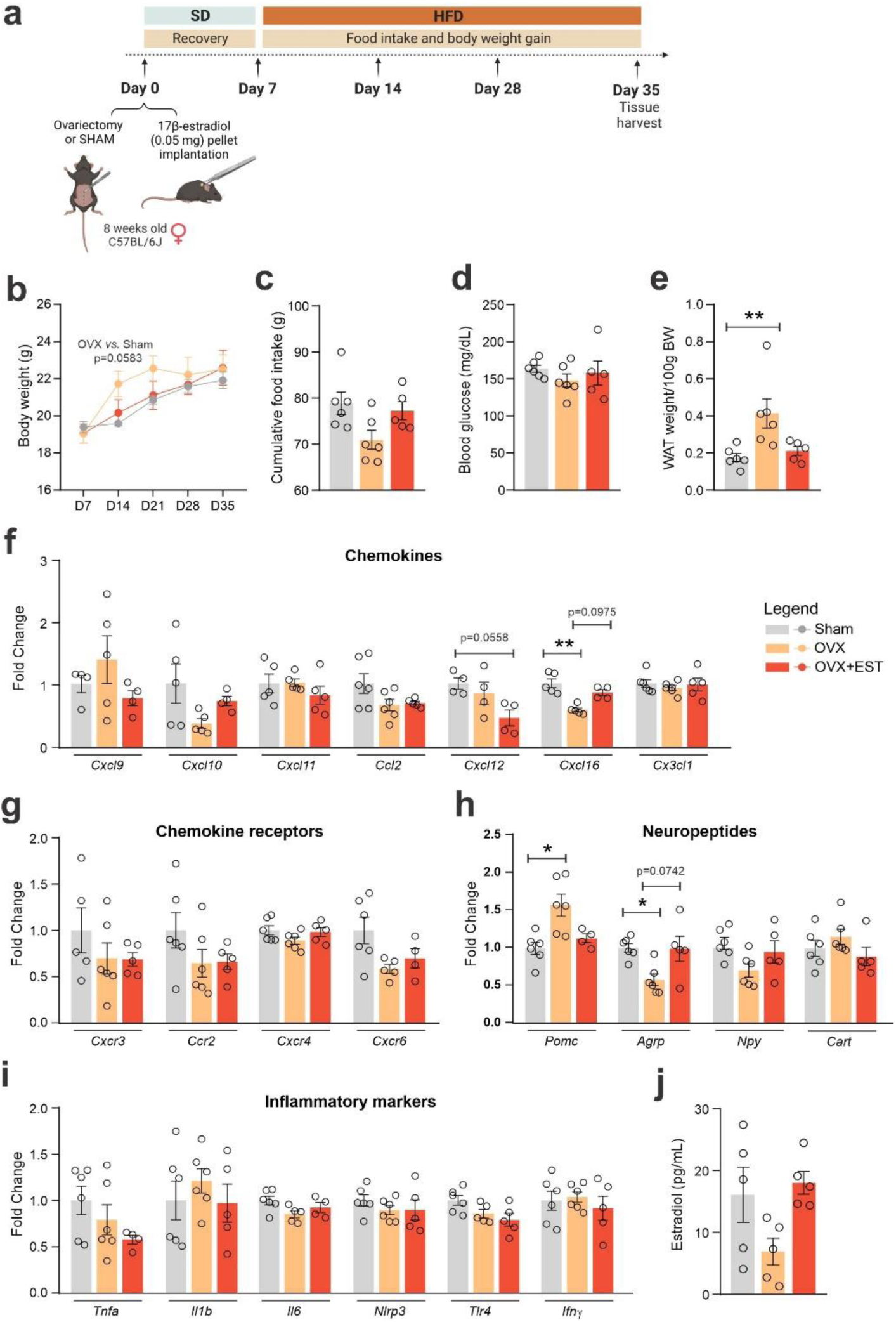
Ovariectomy and ovariectomy with estradiol replacement modulate hypothalamic chemokine and neuropeptides. a) Experimental design of the ovariectomy and estradiol replacement protocol performed in C57BL/6J female mice; b) Body mass gain from Day 7 to Day 35 of the experimental protocol. c) Total food intake during the experimental period. d) Fasting blood glucose levels, and d) Retroperitoneal white adipose tissue depot weight at Day 35. Hypothalamic mRNA levels of f) chemokines; g) chemokine receptors; h) neuropeptides; i) inflammatory genes. Data are expressed as mean ± SEM of 4-6 mice per group. One-way and Two-way ANOVA followed by Sidak’s post-hoc test and Mann-Whitney test were used for statistical analyses. *p<0.05 and **p<001, in comparison with the Sham group.

**Supplementary Figure 2.**
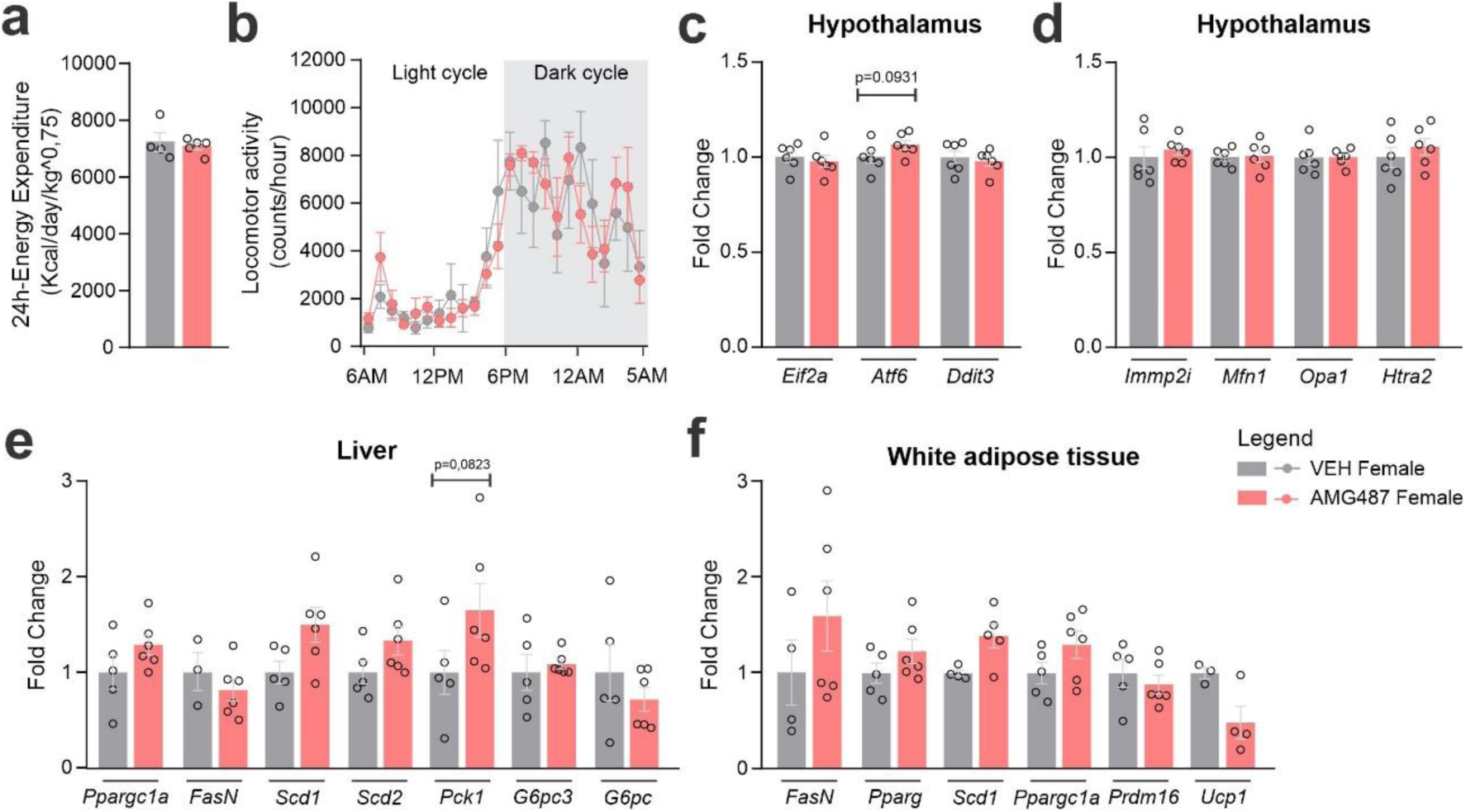
CXCR3 systemic blockage in HFD-fed female mice. a) Total 24h-Energy expenditure and b) Locomotor activity at Day 24 of the experimental protocol. c) Delta body weight during the experimental period. c) Hypothalamic mRNA levels of ER stress genes; d) Hypothalamic mRNA levels of mitochondrial function; e) Hepatic mRNA levels of genes of lipid and glucose metabolism; f) White adipose tissue mRNA levels of genes of lipid metabolism and thermogenesis. Data were expressed as mean ± SEM of 4-6 mice per group. To perform indirect calorimetry and locomotor activity, we have used 5-6 mice/group. To perform qRT-PCR we have used 4-6 mice/group. Two-way ANOVA followed by Sidak’s post-hoc test and Mann-Whitney test were used for statistical analyses.

**Supplementary Figure 3.**
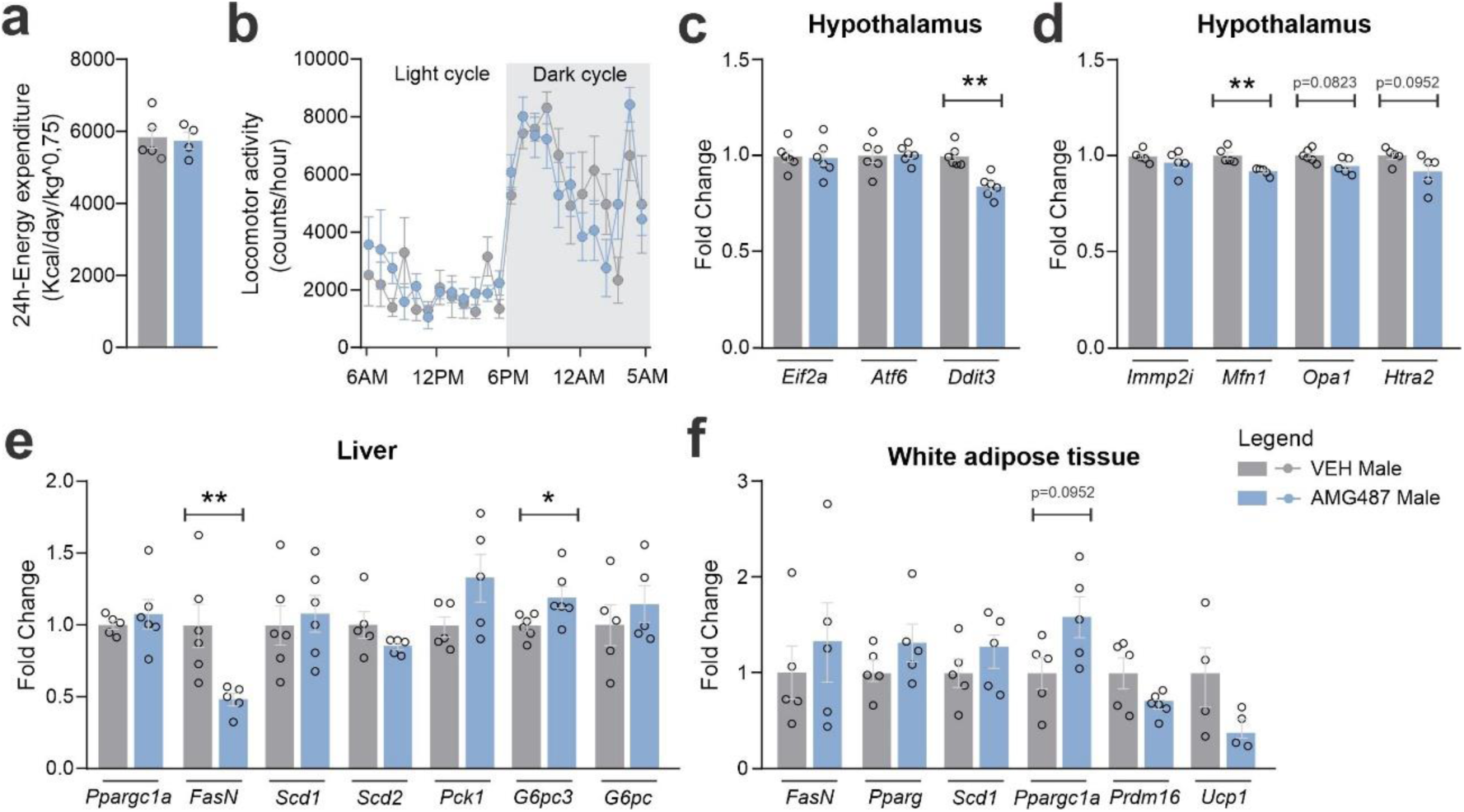
CXCR3 systemic blockage in HFD-fed male mice. a) Total 24h-Energy expenditure and b) Locomotor activity at Day 24 of the experimental protocol. c) Delta body weight during the experimental period. c) Hypothalamic mRNA levels of ER stress genes; d) Hypothalamic mRNA levels of mitochondrial function; e) Hepatic mRNA levels of genes of lipid and glucose metabolism; f) White adipose tissue mRNA levels of genes of lipid metabolism and thermogenesis. Data were expressed as mean ± SEM of 4-6 mice per group. To perform indirect calorimetry and locomotor activity, we have used 5-6 mice/group. To perform qRT-PCR we have used 4-6 mice/group. Two-way ANOVA followed by Sidak’s post-hoc test and Mann-Whitney test were used for statistical analyses. *p<0.05 and **p<001, in comparison with VEH treated group.

## Notes

### Competing Interest Statement

The authors have declared no competing interest.

### Summary of Updates

We included three supplementary figures, that expand the phenotype of the mice under intervention of CXCR3. We also evaluated ovariectomized mice, as the phenotypes are distinct between males and females.

